# Spindle position dictates division site during asymmetric cell division in moss

**DOI:** 10.1101/2020.03.03.975557

**Authors:** Elena Kozgunova, Mari W. Yoshida, Ralf Reski, Gohta Goshima

## Abstract

Asymmetric cell division (ACD) underlies the development of multicellular organisms. It is considered that the division site in land plants is predetermined prior to mitosis and that the localization of the mitotic spindle does not govern the division plane. This contrasts with animal ACD, in which the division site is defined by active spindle-positioning mechanisms. Here, we isolated a hypomorphic mutant of the conserved microtubule-associated protein TPX2 in the moss *Physcomitrium patens* and observed abnormal spindle motility during cell division. This defect compromised the position of the division site and produced inverted daughter cell sizes in the first ACD of gametophore (leafy shoot) development. The phenotype was rescued by restoring endogenous TPX2 function and, unexpectedly, by depolymerizing actin filaments. Thus, we have identified an active spindle-positioning mechanism that, reminiscent of acentrosomal ACD in animals, involves microtubules and actin filaments and sets the division site in plants.

## Introduction

Chromosome segregation during mitosis and meiosis is driven by a complex microtubule (MT)-based apparatus known as the spindle. Animal spindles are known to be mobile and their final position corresponds to the future cytokinesis site, which in turn could determine daughter cell fate after asymmetric division. The mechanism by which the spindle is positioned and spatially controls the assembly of the cytokinetic machinery has been well studied in animals, and the critical roles of force-generating machinery, such as dynamic MTs, actin, and motor proteins, have been elucidated (Bergstralh et al., 2017; Kiyomitsu, 2019). However, in plants, it is believed that the preprophase band (PPB), a plant-specific MT-actin belt formed prior to mitotic entry, determines the future cell division site (Buschmann and Müller, 2019; Rasmussen and Bellinger, 2018; Verma, 2001). The mitotic spindle always forms perpendicular to and at the site of the PPB, perhaps by the action of bridging MTs that connect the spindle to the former PPB site (Ambrose and Cyr, 2008).

The spindle position in plant cells is considered to be static, unless a strong force (1600-3350 × *g*) is applied through centrifugation, which also causes other cytoplasmic components to translocate (Arima et al., 2018; Ôta, 1961). The static nature of spindles is consistent with the fact that plants lack centrosomes, which play key roles in spindle translocation in animal somatic cells (Bergstralh et al., 2017; Kiyomitsu, 2019). Multiple proteins co-localized to the PPB are required to establish and maintain the cortical division zone (CDZ), towards which the cytoskeleton-based cytokinetic machinery, known as the phragmoplast, expands while recruiting cell plate components (Müller, 2019; Smertenko et al., 2017). However, the essential role of PPB in determining the cell division site has been challenged recently by a study that characterized cell divisions in the roots of *Arabidopsis thaliana trm678* mutants lacking PPB. The findings showed that PPB is not essential for division plane determination (Schaefer et al., 2017). In addition, global tissue organization and plant development, although more variable, were comparable to wild-type plants, implying that ACDs are executable without PPBs.

The moss *Physcomitrium patens* (formerly called *Physcomitrella patens*) is an attractive model plant for studying PPB-independent division plane determination, as most cell types naturally lack PPBs, but are capable of oriented cell division and patterning into complex 3D structures, such as gametophores (leafy shoots) (Kosetsu et al., 2017; Moody et al., 2018). We have previously shown that the MT structure, called the gametosome, appears in the cytoplasm transiently at prophase and acts as a determinant of spindle orientation (Kosetsu et al., 2017). However, gametosomes are dispensable for spindle MT generation and spindle positioning.

In animal cells, the mitotic spindle is assembled through rapid MT nucleation and amplification aided by multiple proteins, including γ-tubulin, augmin, and TPX2 (targeting factor for Xklp2) (Petry, 2016). A previous study in *A. thaliana*, using a combination of knockout and cross-species antibody injection, suggested that TPX2 is an essential gene (Vos et al., 2008). However, these results were recently questioned when several viable AtTPX2 t-DNA insertion mutants were obtained (Boruc et al., 2019). In addition to canonical TPX2, several TPX2-like genes lacking one or more functional domains have been identified in *A. thaliana*. Among them, TPX2L3 lacks a C-terminal kinesin-binding motif but is strongly associated with Aurora kinases and is essential for embryogenesis (Boruc et al., 2019). However, the mechanism by which TPX2 contributes to spindle formation and MT amplification in plant cells remains unknown.

In this study, we aimed to characterize TPX2 function in the spindle assembly of *P. patens*, wherein many research tools, including inducible RNAi, endogenous gene tagging, and highly efficient CRISPR are easily applicable (Lopez-Obando et al., 2016; Yamada et al., 2016). In addition to TPX2’s role in MT amplification during early mitosis, we found an unexpected function of TPX2 in maintaining spindle position during asymmetric cell division in gametophores.

## Results

### *P. patens* TPX2

We identified five genes homologous to *TPX2* in the *P. patens* genome using a BLAST search and named them *TPX2-1* to *-5* (Figure 1A). TPX2-1 to -4 proteins showed higher similarity to canonical TPX2 in seed plants (e.g., *A. thaliana* and *Orysa sativa*), whereas TPX2-5 appears to have lost the N-terminal Aurora-binding motif (Boruc et al., 2019; Tomaštíková et al., 2015; Vos et al., 2008), but retained the highly conserved C-terminal domains and, to a certain extent, γ-tubulin activation motifs (Alfaro-Aco et al., 2017) (Figure 1A, B). During mitosis, endogenous TPX2-1, -2, and -4 proteins fused with Citrine (“Cit”) or mNeonGreen (“NG”) or SunTag (“ST”) in-frame at the C- or N-terminus, were localized to the spindle and phragmoplast, with enrichment at the polar region (Figure 1C, Supplemental figure 1, Video 1). A similar localization has been reported for *Arabidopsis* TPX2 (Boruc et al., 2019; Vos et al., 2008). TPX2-5 was observed at the spindle and showed more uniform binding to phragmoplast MTs, suggesting that TPX2-5 might have an additional function (Figure 1C, Supplemental figure 1E-F, Video 1). Unlike TPX2 in animals or seed plants (Boruc et al., 2019; Vos et al., 2008), none of the TPX2 proteins of *P. patens* were sequestered in the nucleus during interphase (Supplemental figure 2).

**Figure 1.**
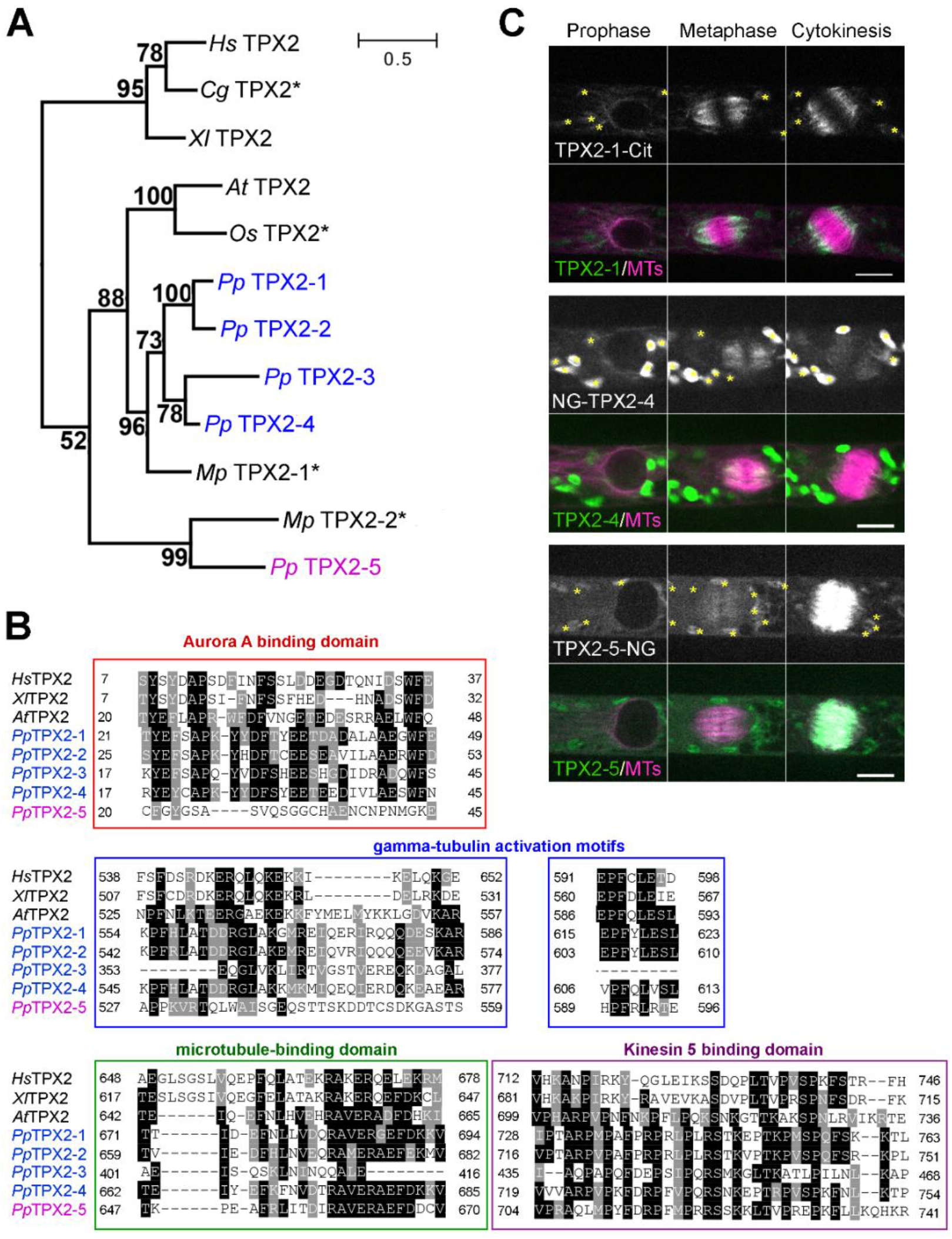
TPX2 homologues and their localization in *P. patens*. **(A)** Phylogeny analysis revealed two distinct groups of TPX2 proteins in *P. patens*: Pp *TPX2-1* to *-4* (blue), which are more similar to *TPX2* genes from seed plants, and atypical *TPX2-5* (magenta). Asterisks mark predicted proteins, numbers show bootstrap values. Bar, 0.5 amino acid substitutions per site. Note that AtTPX2L3 and AtTPX2L2 could not be added to this tree, since they lack the C-terminal region that is conserved in canonical TPX2 proteins. Hs: *Homo sapiens*, Gg: *Gallus gallus*, Xl: *Xenopus laevis*, At: *Arabidopsis thaliana*, Os: *Oryza sativa*, Pp: *Physcomitrium patens*, Mp: *Marchantia polymorpha*. **(B)** Alignment of TPX2 proteins. Conserved residues are boxed, whereas similar amino acids are hatched. **(C)** Localization of endogenous TPX2-1-Citrine, mNeonGreen-TPX2-4 and TPX2-5-mNeonGreen. More uniform spindle localization was detected for TPX2-5. Asterisks indicate autofluorescent chloroplasts. Bar, 10 μm. The full version of mitotic localization data is presented in Supplemental figure 1.

### Generation of hypomorphic mutants of *TPX2-5*

The similarity in amino acid sequences and intracellular localization suggested that TPX2-1 to -4 have redundant functions. Therefore, we simultaneously targeted these genes using a previously established CRISPR/Cas9 protocol (Leong et al., 2018). We isolated a line, named *TPX2 1-4Δ*, in which frameshifts were introduced to all four *TPX2* genes in the exons present in all transcript variants identified in the Phytozome database (Supplemental figure 3A). The *TPX2 1-4Δ* line developed protonema (tissue comprised of frequently dividing tip-growing cells) and gametophores in a similar manner to the parental “GH” line (Figure 2A, Supplemental figure 4). We then attempted to knock out the *TPX2-5* gene in the *TPX2 1-4Δ* background by means of homologous recombination. However, we could not isolate a knockout line after multiple attempts.

**Figure 2.**
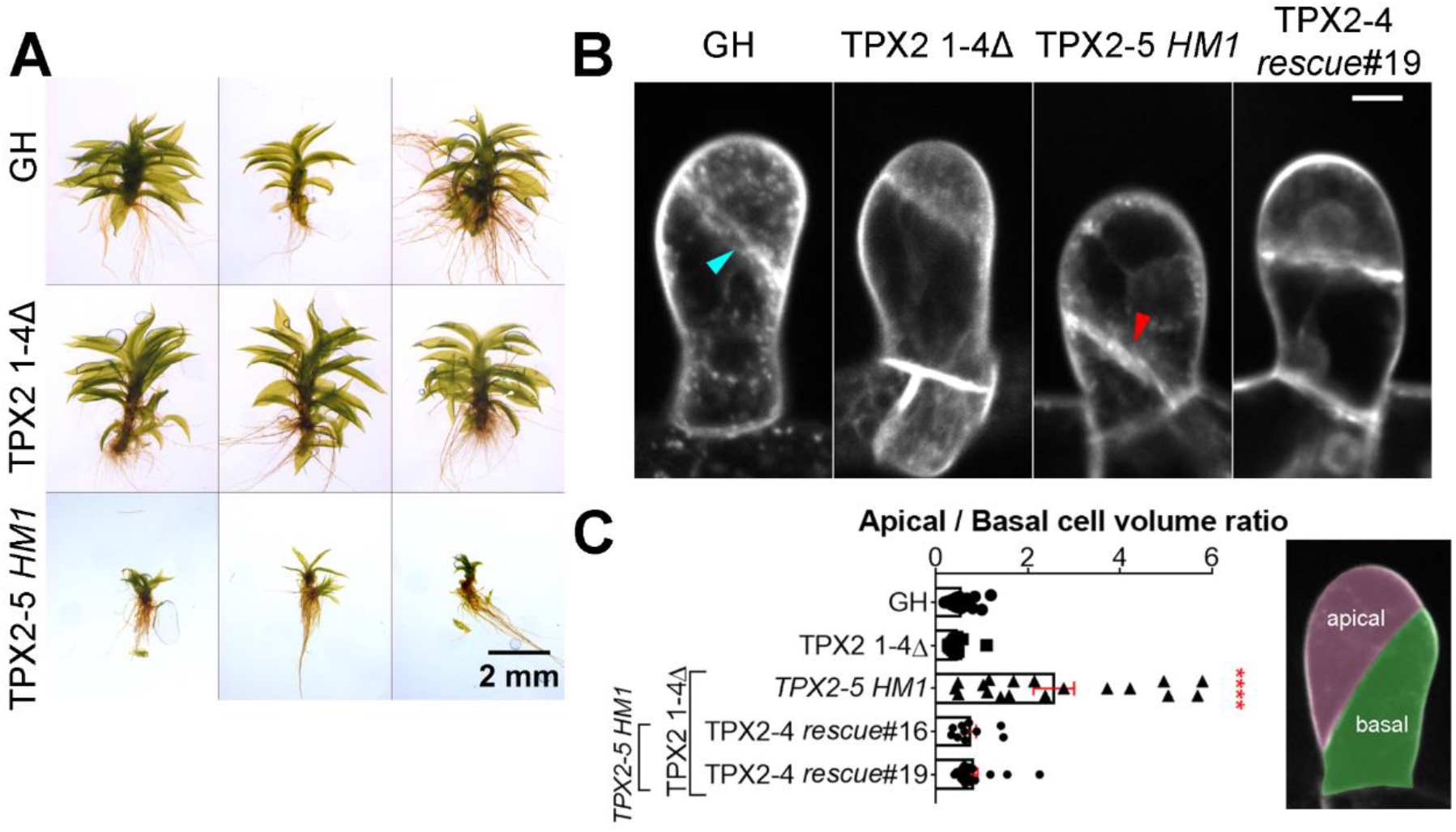
Abnormal cell division site in the gametophore initial of a *TPX2-5 HM1* mutant. **(A)** Representative photos of gametophores after 4 weeks of culture of GH (control), *TPX2 1-4Δ*, and *TPX2-5 HM1* lines. Brightness/contrast have been linearly adjusted. Extended version is presented in Supplemental figure 4. Bar, 2 mm. **(B)** Gametophore initial at the 2-cell stage stained with FM4-64 dye. Normal and defective cell plate positions are indicated with cyan and red arrowheads, respectively. The full version can be found in Figure 2 - figure supplement 4A. Bar, 10 μm **(C)** The apical/basal cell volume ratio was estimated as the apical cell (pink) volume divided by the basal cell (green) volume, measured during the 2-cell stage (mean ± SEM, ****p=0.0001 by one way Anova with Dunnett’s multiple comparison test against GH). n = 22, 15, 18, 10 and 18 for GH, TPX2 1-4Δ, *TPX2-5 HM1*, TPX2-4 *rescue*#16 and TPX2-4 *rescue* #19. The full version can be found in Supplemental figure 6.

Nonetheless, in our attempt to knockout the *TPX2-5* gene, we isolated three lines with defective gametophore development after two independent transformations (Figure 2A, Supplemental figure 4). In these lines, the original *TPX2-5* gene was replaced with a hygromycin cassette, as confirmed by PCR and sequencing (Supplemental figure 3B, C). However, DNA was amplified by PCR using *TPX2-5* “internal” primers, and we confirmed by sequencing that all exons of the *TPX2-5* gene remained (Supplemental figure 3D). These data suggested that the *TPX2-5* gene removed from the original locus was re-inserted into another genomic locus, possibly through micro-homology recombination. In all three mutants we isolated, the expression of *TPX2-5* mRNA was compromised (Supplemental figure 3E). Hereafter, these <u>h</u>ypo<u>m</u>orphic lines are referred to as *“TPX2-5 HM1,” “TPX2-5 HM2”* and *“TPX2-5 HM3.”*

To test possible redundancy between *TPX2-5* and other *TPX2s*, we performed a rescue experiment in which full-length *TPX2-4* (with Cerulean tag) at the native locus was expressed in the *TPX2-5 HM1* line (Supplemental figure 5A, E). The lines no longer showed a mitotic delay or dwarf gametophore phenotype (Supplemental figure 4B, Supplemental figure 5B-D). In addition, we attempted to knockout the *TPX2-5* gene in the wild-type background (i.e., other *TPX2* genes were intact). Once again, we could not isolate a real knockout line, but isolated a *TPX2-5* translocation line termed *TPX2-5 M* (Supplemental figure 3). Given the high efficiency of homologous recombination in *P. patens*, the *TPX2-5* gene is likely essential. However, in the wild-type background, relocation of the *TPX2-5* did not yield any noticeable phenotypes, consistent with the result of the rescue experiment (Supplemental figure 4). These data suggest that *TPX2-5* has redundant as well as distinct functions compared to the other TPX2s.

### Spindle motility and division plane shift in the *TPX2* mutant gametophore

Next, we aimed to determine the role of TPX2 in the mitosis of gametophore cells, as dwarf gametophores are the most prominent phenotype of the *TPX2-5 HM1,2,3* lines (Figure 2A, Supplemental figure 4B). Dwarf organ development in plants is sometimes associated with defective cytokinesis (Martinez et al., 2017). Thus, we first used the lipophilic dye FM4-64 to visualize cell plates in gametophore initials (stem cells) after the first cell division. We observed that the position of the cell plate shifted to the basal side of the gametophore initial in multiple cells in all three *TPX2-5* mutants, dramatically skewing the size ratio between apical and basal daughter cells (Figure 2B, C, Supplemental figure 6, Video 2). This was due to an overall reduced level of TPX2, as the cell plate positioning was normal in the aforementioned *TPX2-4* rescue lines and *TPX2-5 M* line (Figure 2B, C, Supplemental figure 6, Video 2). To investigate what caused defects in the cell division site in the *TPX2-5 HM1,2,3* lines, we next performed live-cell imaging of the first cell division in the gametophore initial.

To observe mitotic MTs in gametophore initial cells, we introduced another marker, mCherry-tubulin, to *TPX2-5 HM1, TPX2 1-4Δ*, and control GH lines (note that histone and tubulin were labeled with the same color). The majority of gametophore initial cells in the *TPX2-5 HM1* line formed a bipolar spindle (90%; *n* = 19; Figure 3A, B), although slightly smaller than the spindle in the control or *TPX2 1-4Δ* lines (Figure 3C). Strikingly, we observed that the bipolar metaphase spindle moved unidirectionally over 5 μm to the basal side in 73% (n = 14) of *TPX2-5 HM1* cells (Figure 3A, D, Video 3). Consequently, the phragmoplast formed close to the basal edge. We concluded that defective spindle positioning after NEBD causes abnormal cell plate position in the gametophore initials of the *TPX2-5 HM1* line.

**Figure 3.**
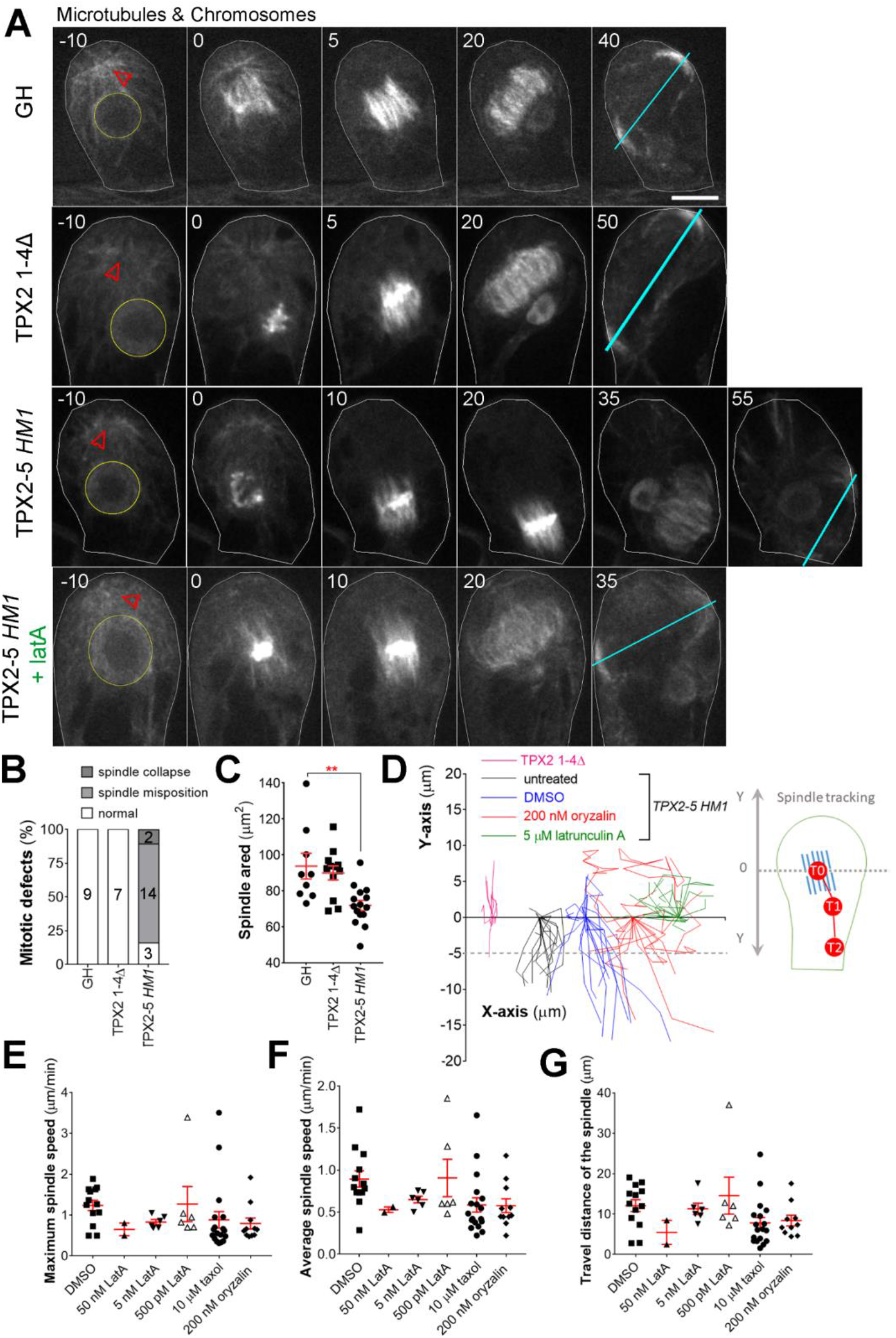
Spindle position is actively maintained through the interplay between microtubules and F-actin. **(A)** Live-cell imaging of the first asymmetric division in the gametophore initial revealed a link between the metaphase spindle and phragmoplast positioning. The positions of the nucleus and gametosome (prophase MTOC appeared in the apical cytoplasm) are indicated with yellow circles and red arrowheads, respectively. Cyan lines show the position and orientation of the phragmoplast. Cell borders are outlined with white lines. Bar, 10 μm. **(B)** The frequency and type of spindle defects in gametophore initial mitosis observed in GH (control), *TPX2 1-4Δ*, and *TPX2-5 HM* lines. Numbers within the columns indicate number of cells with corresponding phenotypes. **(C)** Area occupied by the metaphase spindle (spindle size) in gametophore initials. (mean ± SEM; \*\**p* = 0.0029, two-tailed Student’s t-test) **(D)** Tracking of the spindle center position from NEBD to anaphase onset. We assigned the starting position as Y = 0 and different X positions for each sample group. Note that after 5 μM latrunculin A treatment, spindles never showed motility towards the basal end of the cell, i.e. negative Y-values. Each line represents spindle movement in a single cell. More than 12 cells were observed for each sample group in three or more independent experiments. **(E)** Maximum spindle speed (μm/min), **(F)** average spindle speed (μm/min), and **(G)** distance travelled by the spindle (μm) in the *TPX2-5 HM1* cell line under various treatments. Total number of cells observed, including those without spindle motility: n = 20 (DMSO), 18 (50 nM LatA), 12 (5 nM LatA), 13 (500 pM LatA), 23 (10 μM taxol) and 16 (200 nM oryzalin). Only spindle motility towards the basal end of the cell was analyzed. Bars represent mean ± SEM

In six cases, spindle motility or rotation was observed in the second or later divisions, while the first cell division site appeared normal (Video 4). This suggested that defective ACDs in multiple stages underlie dwarf gametophore formation. Defects in cell expansion could be another possible explanation for the dwarf gametophore phenotype, as gametophore cells grow by expansion, controlled by turgor pressure and cytoskeleton dynamics. To test this possibility, we analyzed cell expansion rates in early gametophore development (after the first cell division) in the control (GH) and *TPX2-5 HM1* lines (Supplemental figure 7A). We did not detect a statistically significant difference in the increase in cell size during one cell cycle (Supplemental figure 7B). Thus, in the early stages, spindle mispositioning appears to be a primary cause of defective gametophore development in the *TPX2-5 HM1* line.

### Spindle motility in the *TPX2* mutant is actin-dependent

In animals, spindle motility relies on MTs and actin. To test the involvement of the cytoskeleton in spindle motility, we first partially depolymerized MTs in the *TPX2-5 HM* line using low-dosage oryzalin (200 nM), a MT-destabilizing drug. Upon treatment, the spindle behaviors were more variable and showed overall differences from the untreated cells. Most notably, in 5 of 16 cells, we observed that the spindle had shifted towards the apical side of the cells, which was not observed in untreated cells (Figure 3D). In addition, the maximum spindle speed was decreased (Figure 3E). In the presence of oryzalin, 9 out of 11 spindles moved slower than 1 μm/min (maximum speed), whereas only 2 out of 13 did so in control cells (DMSO). These data suggest that MTs contribute to spindle motility to a certain extent.

We also tested the effect on spindle motility of the MT-stabilizing drug taxol at a final concentration of 10 μM. The frequency of cells showing spindle motility did not decrease (18 of 23 cells). However, the motility was slower and the traveling distance was shorter than that of the control cells (Figure 3E-G). The difference was not statistically significant, possibly because the spindle motility phenotype showed high variability in the *TPX2-5 HM1* line. This positive but mild effect might be due to the low sensitivity of *P. patens* to taxol (Doonan et al., 1988). For example, we never observed mitotic arrest after taxol treatment, which is expected after high-dose taxol treatment of animal cells (Jordan et al., 1993).

In early mitosis in the gametophore initial cell, actin is broadly distributed in the cytoplasm, including the spindle area, and actin depolymerization by latrunculin does not affect spindle morphology, orientation, or positioning in wild-type (Kosetsu et al., 2017). We investigated the effect of latrunculin on spindle motility of the *TPX2-5 HM1* mutant cells. Interestingly, 5 μM latrunculin A treatment completely (20 of 20 cells) suppressed the basal motility of the spindle in the *TPX2-5 HM1* mutant line (Figure 3A, D, Video 3). In addition, we tested several lower concentrations of latrunculin A and found a concentration-dependent response in spindle motility. For example, only 2 of 18 gametophore initials still displayed basal spindle motility when treated with 50 nM latrunculin A (Figure 3E-G). We also attempted to visualize actin distribution during spindle movement. To this end, we introduced mCherry-Lifeact (actin marker) in the *TPX2-5 HM1* background. However, we observed that introducing mCherry-Lifeact rescues the spindle motility in all three lines independently selected (no motility was observed in 15 cells). It is known that Lifeact expression can alter actin dynamics and organization in diverse cell types (Flores et al., 2019; Courtemanche et al., 2016, Spracklen et al., 2014), likely by competing for binding sites with actin-interacting proteins (Belyy et al., 2020).

We speculate that this was also the case in *P. patens*. We concluded that actin is a main driving force of basal spindle motility in the mutant.

### Roles of TPX2-5 in spindle assembly in protonema

ACD occurs in gametophores and in protonema tissue. The protonema consists of tip-growing chloronemal and caulonemal cells. The apical, tip-growing cells frequently divide, while subapical cells occasionally form a bulge and divide asymmetrically, producing a new apical tip cell, in a process known as “branching”. We did not observe spindle motility in the protonema of the *TPX2-5 HM1* line (n = 85, including caulonemal/chloronemal apical cells and branching cells). The identified differences from the wild type were the cell plate orientation and apical cell growth speed in caulonemal cells, which might reflect a slight alteration in MT dynamics in the mutant (Supplemental figure 8). However, it is unlikely that these protonema defects have a significant impact on gametophore development.

A rare spindle phenotype observed in both protonema (n = 1 of 85) and gametophore initial cells (n = 2 of 19) was spindle collapse, which led to chromosome missegregation, cytokinesis failure, and multinucleation (Figure 3B, Video 5). In an earlier study, we showed that a certain percentage of protonema cells of *P. patens* were able to recover after cytokinesis failure and re-enter the cell cycle, resulting in whole genome duplication (Kozgunova et al., 2019). In the context of gametophore development, diploid *P. patens* has fewer gametophores for unknown reasons (Schween et al., 2005). To exclude the possibility that *HM* mutants have a high number of diploid protonema cells, compromising gametophore development, we conducted flow cytometry analysis of ploidy levels in the protonema cells of control (GH), *TPX2 1-4Δ*, and *TPX2-5 HM1,2,3* lines. We detected no additional peaks that would indicate higher ploidy in any of these lines (Supplemental figure 9).

To gain insights into the essential mitotic function of TPX2-5, we performed a detailed analysis of cell division phenotypes in the mutant with the greatest effect. To this end, we selected an inducible *TPX2-5* RNAi lines in the *TPX2 1-4Δ* background. Since RNAi induction almost completely inhibited cell growth and gametophore development, we focused on the division of the protonemal apical stem cells that appear earlier than gametophores. Using time-lapse imaging of MTs and chromosomes, we observed severe MT phenotypes during the early stages of mitosis (Figure 4). During prophase, we detected a reduction in the number of perinuclear MTs (Figure 4E), which was also observed with a γ-tubulin RNAi (Nakaoka et al., 2012), and abnormal nuclear shape, which is unique to the *TPX2 1-4Δ* mutant (elongated nucleus prior to NEBD, Figure 4C). After NEBD, 3 out of 52 cells in the RNAi lines failed to form bipolar spindles, followed by metaphase arrest and chromosome missegregation, which was never observed in the control lines (Figure 4D). Other cells formed bipolar spindles; however, the number of MTs in the prometaphase spindle was greatly reduced (Figure 4F, blue). This was no longer the case at metaphase, where the number of MTs was similar to control spindles, indicating the recovery of MT numbers during prometaphase (Figure 4F, green). Phragmoplast formation and expansion were similar to those in control cells. This is opposite to the findings in augmin-knockdown cells, in which the number of metaphase spindle MTs and phragmoplast MTs, but not prophase MTs, was reduced (Nakaoka et al., 2012). A plausible interpretation is that MTs were generated and reached control levels through γ-tubulin and augmin-dependent MT amplification during prolonged prometaphase in the *TPX2* lines (Petry et al., 2013). Thus, our results suggest that the role of TPX2 in MT amplification is dominant during early mitosis. As γ-tubulin activation motifs (Alfaro-Aco et al., 2017) are partially conserved in all moss TPX2 proteins (Figure 1B), we suggest that TPX2 is required for γ-tubulin-dependent MT nucleation in prophase and prometaphase, while augmin takes over from prometaphase. Other phenotypes observed upon TPX2-5 depletion included chromosome missegregation (29%, n = 15) and spindle misposition/orientation (33%, n = 17, Figure 4D, Video 6). Overall, these phenotypes suggested the functional conservation of moss TPX2 with well-studied animal orthologues, namely assisting in MT formation through nucleation and/or stabilization.

**Figure 4.**
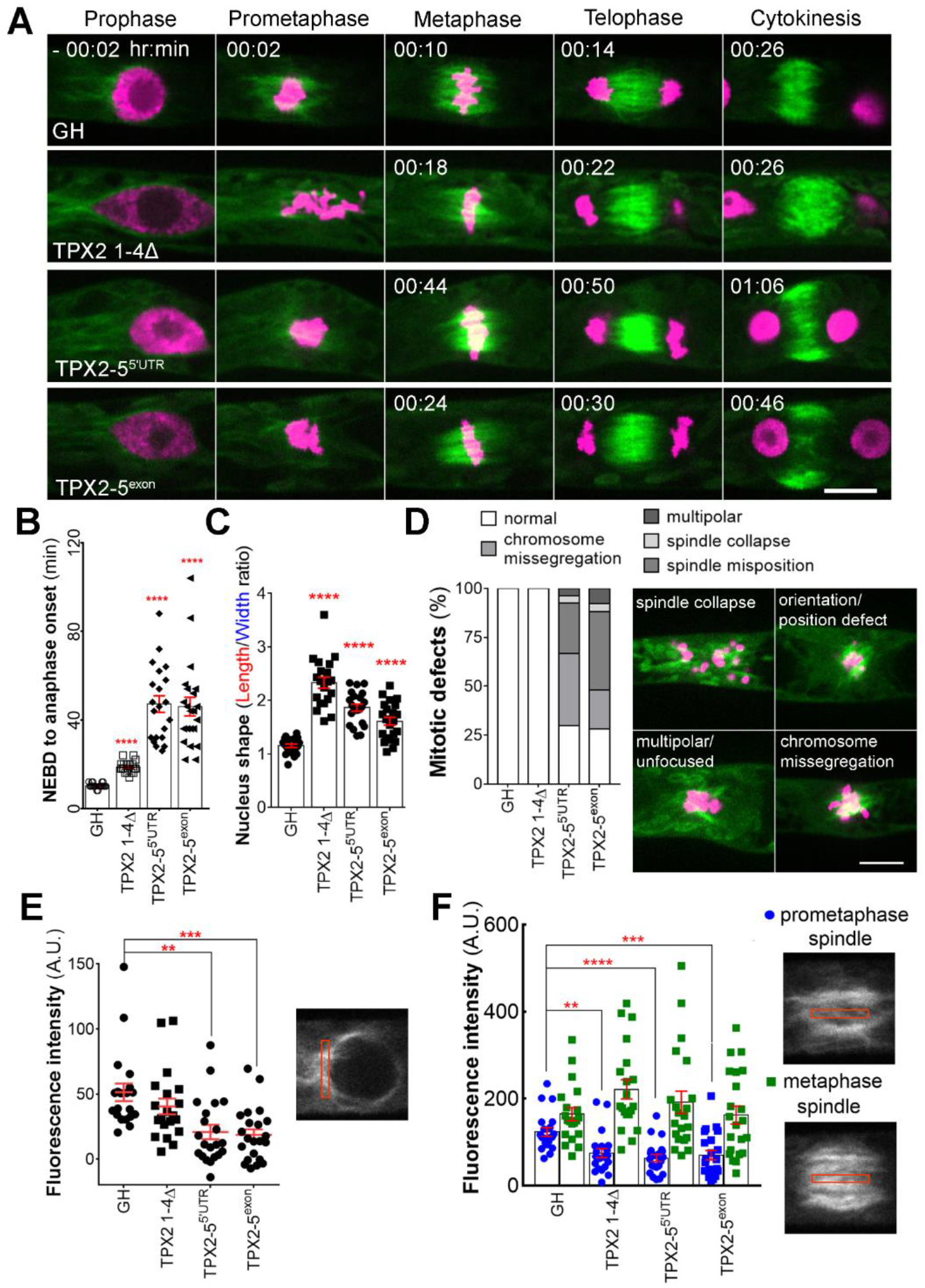
TPX2 contributed to microtubule amplification in early mitosis. **(A)** Representative images of the mitosis of protonemal apical cells in GH (control), *TPX2 1-4Δ, TPX2-5^5’UTR^* RNAi, and *TPX2-5^exon^* RNAi lines. Green, GFP-tubulin; Magenta, histoneH2B-mRFP. Bar, 10 μm. **(B)** Mitotic duration calculated from NEBD to anaphase onset (mean ± SEM; \*\*\*\**p* < 0.0001, two-tailed Student’s t-test). **(C)** Nucleus shape prior to NEBD, measured as a ratio of nucleus length to nucleus width (mean ± SEM; ****p < 0.0001, two-tailed Student’s t-test). **(D)** Frequency and type of mitotic defects observed. Bar, 10 μm. **(E)** Fluorescence intensity of perinuclear MTs (mean ± SEM; \*\**p* = 0.0011, \*\*\**p* = 0.0002; two-tailed Student’s t-test). A.U. stands for Arbitrary Units. **(F)** Fluorescence intensity of MTs in the prometaphase spindle (4 min after NEBD) and metaphase spindle (2 min before anaphase onset), measured from a single focal plane, with the cytoplasmic background subtracted. A decrease in fluorescence intensity was detected in prometaphase, but not in metaphase spindles (mean ± SEM \*\**p* = 0.0018, \*\*\*\**p* ≤ 0.0001, \*\*\**p* = 0.0008; two-tailed Student’s t-test).

## Discussion

In plants, defective division sites have been mostly attributed to defects in PPB formation (Schaefer et al., 2017; Yoneda et al., 2005), phragmoplast guidance errors (CDZ deficiency) (Lipka et al., 2014; Müller, 2019), or abnormal positioning of the nucleus in prophase (Kimata et al., 2019; Yamada and Goshima, 2018). The current study identified an independent and hitherto unappreciated cause of division site abnormality: spindle motility after NEBD. Furthermore, we identified a conserved protein, TPX2, which plays a critical role in spindle positioning in *P. patens*. Interestingly, a recent study using neural stem cells of the embryonic developing mouse neocortex showed that TPX2 knockdown not only affects spindle MT generation but also spindle orientation, implicating a conserved TPX2-dependent mechanism of spindle positioning (Vargas-Hurtado et al., 2019). Spindle-specific positioning defects in meiosis II have also been observed in *jas* and *ps1* mutants of *Arabidopsis* due to abnormal organelle distribution (Brownfield et al., 2015). However, meiosis II is a unique system in which two spindles share a cytoplasm and may be partially fused in the absence of an organelle barrier. Therefore, there is unlikely to be a mechanistic analogy to moss gametophores. Our data suggest that defective ACD underlie dwarf gametophores in the *TPX2* mutant. However, it cannot be ruled out that non-mitotic functions of TPX2-5 are also required for proper gametophore development. During gametophore differentiation, MT organization undergoes changes to cortical arrays and PPB-like structure has been observed in mature gametophores (Doonan et al., 1987; Kosetsu et al., 2017). An interesting future study would be to investigate whether moss TPX2-5, which is not sequestered in the nucleus unlike animal TPX2, regulates interphase MT dynamics and PPB formation in mature gametophore cells.

In addition to TPX2, this study uncovered the involvement of actin in spindle positioning. This was also an unexpected observation, as the function of actin in plant cell division has been mostly attributed to phragmoplast guidance during cytokinesis (Livanos and Müller, 2019; Rasmussen and Bellinger, 2018). Our data suggest that actin filaments are the major driving force in basal spindle motility, while MTs counteract this force under normal conditions. When the overall levels of TPX2 are decreased, fewer and/or less stable MTs are insufficient to hold the spindle in place (Figure 5). This model is supported by the observation that 5 μM latrunculin A completely abolished basal spindle motility and that a less severe spindle motility phenotype was observed after taxol treatment, which partially stabilized MTs.

**Figure 5.**
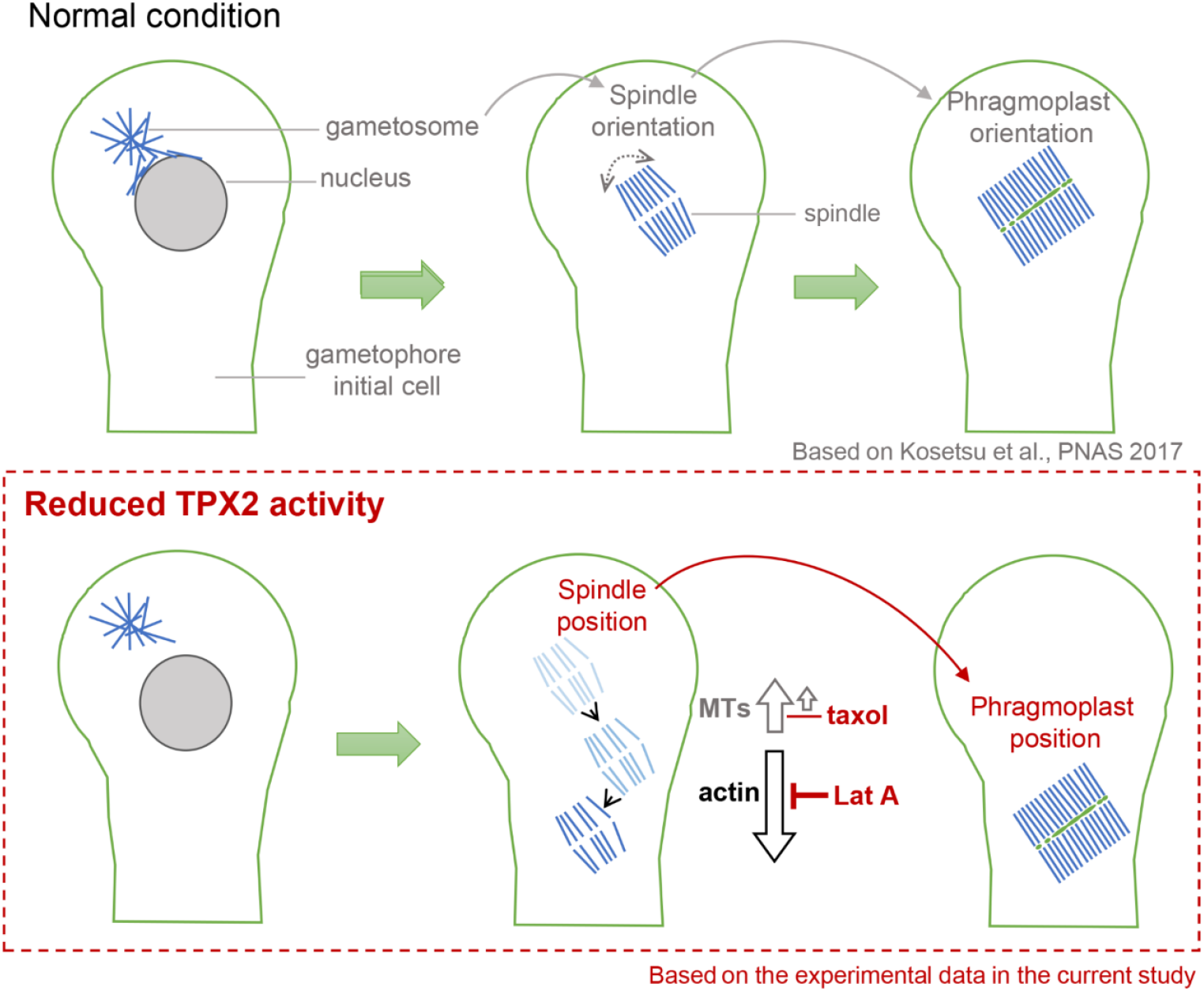
Schematic representation of the spindle impact on phragmoplast position and orientation in the gametophore initial cells of *P. patens*. The upper panel shows the role of the microtubule-organizing center (gametosome) in spindle and phagmoplast orientation, based on Kosetsu et al. (2017). The bottom panel summarizes the findings of the present study and the role of the microtubule-associated protein TPX2 in spindle positioning. In the scenario where TPX2 function is reduced, the spindle can be transported to the bottom of the gametophore initial cell, compromising the cell plate position and daughter cell ratios in asymmetric cell division. This spindle transport is an actin-dependent process, as it can be completely inhibited by depolymerizing actin filaments with latranculin A (Lat A). Furthermore, stabilizing microtubules with taxol could partially counteract spindle transport, suggesting that under normal conditions, microtubules are able to fix spindle position against actin force.

Actin is known to play an important role in spindle positioning in animal cells (Almonacid et al., 2014). Of particular interest are animal oocytes, as they lack centrosomes. In mouse oocytes, spindle migration and symmetry breaking are driven by changes in the stability of the actin meshwork, the formation of which depends on actin nucleators, such as formin-2 and myosin II motor (Almonacid et al., 2014; Duan et al., 2020). Thus, an analogous mechanism may transmit force to transport spindles in moss. However, it should be noted that actin function emerged only in the background of a *TPX2* mutation. Thus, sufficient numbers of spindle MTs may predominate in the “tug-of-war” against actin-dependent forces in wild-type moss cells. TPX2 may be a promising candidate for investigations to elucidate the PPB-independent mechanism of cell division site positioning in seed plants.

## Materials and methods

### *P. patens* culture and transformation

*P. patens* culture and transformation protocols have been described in detail elsewhere (Yamada et al., 2016). In brief, BCDAT agar medium (BCDAT stands for stock solutions <u>B</u>, <u>C</u>, <u>D</u>, and <u>a</u>mmonium <u>t</u>artrate) was used for regular culturing at 25°C under continuous light. Transformation was performed using a standard polyethylene glycol-mediated method using protoplasts. Transgenic lines were selected with the corresponding antibiotics and confirmed by genotyping PCR or sequencing in the case of CRISPR-generated lines. The GH line, expressing histone H2B-mRFP and GFP-α-tubulin, was used for transformation in the CRISPR and knockout experiments, while the mCherry-α-tubulin #52 line was used for Citrine and mNeonGreen endogenous tagging. The transgenic lines generated in this study are listed in the Supplemental Table 1.

### Molecular cloning

Plasmids for Citrine or mNeon-Green endogenous tagging were assembled using In-Fusion (Clontech, Mountain View, CA, USA), in which Citrine or mNeonGreen genes, a G418 resistance cassette (only C-terminal tagging), and homologous recombination regions (500-800 bp of the respective genes) were connected. A similar strategy was used to assemble the knockout plasmid for *TPX2-5*, wherein a hygromycin resistance cassette was flanked by the 5’ and 3’UTR regions of the *TPX2-5* gene. A detailed protocol for endogenous gene tagging and knockouts in *P. patens* has been previously published (Yamada et al., 2016). CRISPR gRNAs targeting one of the exons were designed using the online tool, CRISPOR (http://crispor.tefor.net/), based on target gene specificity (off-target score) and predicted frameshift efficiency. Individual gRNAs were ligated into pCasGuide/pUC18 vector pre-digested with BsaI. Next, gRNA sites, together with the *U6* promoter and gRNA scaffold region, were amplified by PCR and assembled into a single multi-gRNA plasmid, also containing a hygromycin resistance cassette for transient plant selection. The CRISPR protocol has been described in detail elsewhere (Leong et al., 2018). RNAi vectors were cloned using the Gateway system (Invitrogen, Carlsbad, CA, USA), with pGG624 as the destination vector. Two independent, non-overlapping RNAi constructs were prepared for each gene. The full list of plasmids and primers used in this study are shown in Supplemental Table 2.

### Gene expression analysis with quantitative real-time PCR (qRT-PCR)

To determine the expression levels of *TPX2-5* and *TPX2-4* genes, total RNA was extracted from 7-day-old protonema tissue using the innuPREP Plant RNA kit (Analytik Jena GmbH, Berlin, Germany) according to the manufacturer’s instructions. Total RNA was treated with DNase I (Thermo Fisher Scientific) at 37°C for 1 h to remove genomic DNA contamination, and RNA quality was checked by gel electrophoresis. For cDNA synthesis, we used TaqMan Reverse Transcription Reagents (Thermo Fisher Scientific) with random hexamers, according to the manufacturer’s protocols. Primers were designed using the Universal Probe Library (Roche, https://lifescience.roche.com/en_de/brands/universal-probe-library.html#assay-design-center) and selected according to the lowest amount of off-target hits identified in the *P. patens* transcriptome in Phytozome (www.phytozome.net). The efficiency of each primer pair was confirmed prior to analysis with a qPCR run using a series of cDNA dilutions (1:2) and a control without cDNA. Melting curves were analyzed to exclude primer pairs with off-target amplification. The primer pairs used for qRT-PCR are listed in Supplemental Table 2. Samples using a cDNA equivalent to 50 ng of total RNA from corresponding lines were prepared in triplicate or, alternatively, two sets of triplicates for TPX2-5 in the *TPX2-5 HM1, HM2*, and *HM3* lines, with the SensiMix Kit and SYBR Green (Bioline, Luckenwalde, Germany). Negative controls without template addition, as well as reverse transcription controls (reverse transcription reaction carried out without Multiscribe™ RT), were performed for each primer pair and each line. Amplification was performed in 40 cycles at a melting temperature of 60°C. qRT-PCR data were analyzed using LightCycler R 480 software (Roche). The relative expression levels of the gene of interest (GOI), TPX2-5, and TPX2-4, were compared to those of the control (ctrl), internal housekeeping genes EF1α (Pp3c2_10310V3.1), and the ribosomal protein L21 (Pp3c13_2360V3.1), and was calculated as 2^(-ΔCT)^, wherein ΔC_T_=C_T_(GOI)-C_T_(ctrl) and primer efficiency was assumed to be 2. C_T_ was defined as the cycle number at which each sample reached an arbitrary threshold (Livak and Schmittgen, 2001).

### Sample preparation for live-cell imaging

The sample preparation method for live cell imaging has been described in detail in a previous study (Yamada et al., 2016). In brief, a glass-bottom dish coated with a thin layer of BCD agar medium (BCD stands for stock solutions <u>B</u>, <u>C</u>, and <u>D</u>. Details can be found in (Yamada et al., 2016)), was inoculated with moss protonema and cultured under continuous light at 25°C prior to observation. For gametophore induction, 1 μM of the synthetic cytokinin, benzylaminopurine, or 1μM 2iP (2-isopentenyladenine), diluted in 1 mL of distilled water, was added to the 6 – 7-day-old colony and incubated for 10 min. Next, the remaining liquid was aspirated with a pipette, the dish was sealed, and the sample was cultured as described above for 20 – 24 h prior to gametophore imaging. Latrunculin A, taxol or oryzalin were diluted in 1 mL of distilled water to final concentrations. Prior to drug treatment, most of the agar pad from the glass-bottom dish was cut and removed to minimize dilution. For RNAi induction, 400 μL of 5 μM β-estradiol, diluted in distilled water, was added to the pre-cultured protonema 4 d prior to observation. Although β-estradiol was previously supplemented directly to the agar medium (Miki et al., 2016), we found that it almost entirely inhibited cell growth in *TPX2-5* RNAi lines; hence, the protocol was modified.

### Microscopy and data analysis

Sample preparation is described above. Localization, RNAi, and most of the gametophore images were acquired using a Nikon Ti microscope (60 × 1.30-NA lens; Nikon, Tokyo, Japan) equipped with a CSU-X1 spinning-disk confocal unit (Yokogawa, Tokyo, Japan) and an electron-multiplying charge-coupled device camera (ImagEM; Hamamatsu, Hamamatsu, Japan). The microscope was controlled using the NIS-Elements software (Nikon). Imaging after FM4-64 staining, 10 μM taxol treatment, and low-dosage latrunculin A treatment (50 nM, 5 nM, and 500 pM) was performed using a ZEISS inverted microscope (25 × 0.8-NA lens or 63 × 1.4-NA lens, Carl Zeiss Microscopy, Germany) equipped with a CSU-X1 spinning-disk confocal unit (Yokogawa, Tokyo, Japan) and CCD camera (Photometrics Prime sCMOS). All imaging was performed at 22 – 24 °C in the dark, except for the first division of the gametophore initial, since gametophore development requires light (3-min light/2-min dark cycle). For single-leaf imaging, we dissected gametophores using syringe needles and scissors to isolate single leaves. Leaves were mounted in a drop of water between two coverslips, and images were acquired with a Nikon Ti microscope (10 × 0.30-NA lens) in bright-field mode. and image data were analyzed using ImageJ software (National Institutes of Health, Bethesda, MD, USA). Prism software was used to plot the graphs and perform statistical analyses (GraphPad, San Diego, CA, USA). Gametophore and moss colony images were acquired after 4 weeks of culture using a stereoscopic microscope (SZ2-ILST, Olympus Corporation, Tokyo, Japan) equipped with a digital camera (Axiocam ICc1; Carl Zeiss Microscopy, Germany) or a Canon EOS 400 camera, respectively.

### Cell volume analysis

Cell volume was analyzed in gametophore initial cells stained with FM4-64 dye diluted in distilled water to a final concentration of 10 μM. The dye was added to the live-imaging dish before acquisition, without cutting the agar. Imaging was performed immediately after FM4-64 application. Images were acquired as a Z-stack (40 μm, 0.97 μm step) using a ZEISS inverted microscope (25 × 0.8-NA lens) equipped with a CSU-X1 spinning-disk confocal unit (Yokogawa, Tokyo, Japan) and a CCD camera (Photometrics Prime sCMOS). Cell boundary predictions and segmentation were performed using PlantSeg software (Wolny et al., 2020). Default conditions of PlantSeg were used with the following modifications: images were rescaled using in-build function (for our images voxel size was 0.26 μm on the X-Y dimensions and 0.97 μm on the Z dimension) and cell minimum size was set at 400 000 voxels. The accuracy of the cell predictions was confirmed by comparing PlantSeg output files to raw images for each cell. In some cases, when cell boundaries were not predicted accurately, images were deconvolved using Huygens Professional version 19.04 (Scientific Volume Imaging, Hilversum, Netherlands) and again run through PlantSeg with a minimum cell size of 300 000 voxels. If this did not improve the cell boundary predictions, images were excluded from the analysis. Cell volume was extracted from PlantSeg output tiff files using a threshold function and the Voxel counter plugin in ImageJ. 3D projections in Video 2 were created with Imaris software (Bitplane) version 9.7.0.

### Flow cytometry analysis of ploidy

Moss protonema was homogenized in 5 mL of distilled water and cultured on BCDAT agar plates covered with cellophane for 6 days before measurement. Next, small portion of protonema was chopped with a blade in 4’, 6-diamidino-2-phenylindole (DAPI) solution (0.01 mg/L DAPI, 1.07 g/L MgCl_2_·6H_2_O, 5 g/L NaCl, 21.11 g/L TRIS in 1 ml Triton) and filtered through a 30 μm sieve. Fluorescence of the nuclei was determined with a PAS cell analyzer (Partec, Munster, Germany) using a 100 W high-pressure mercury lamp (Schween et al., 2003).

### Sequence analysis

We used MAFFT ver. 7.043 (https://mafft.cbrc.jp/alignment/software/) to align the amino acid sequences of the selected full-length proteins, and then manually revised them with MacClade ver. 4.08 OSX (www.macclade.org) to remove gaps. The Jones-Taylor-Thornton (JTT) model was used to construct maximum-likelihood trees using MEGA5 software (www.megasoftware.net). Statistical support for internal branches by bootstrap analysis was obtained using 1,000 replications. The gene sequence information discussed in this article is available under the following accession numbers in Phytozome (www.phytozome.net): *At*TPX2 (AT1G03780.3); *Os*TPX2 (LOC_Os07g32390.1); *Pp*TPX2-1 (Pp3c17_11160V3.1); *Pp*TPX2-2 (Pp3c1_25950V3.1); *Pp*TPX2-3 (Pp3c24_8590V3.2); *Pp*TPX2-4 (Pp3c23_4540V3.1); *Pp*TPX2-5 (Pp3c5_10270V3.1); *Mp*TPX2-1 (Mapoly0016s0083.1); *Mp*TPX2-2 (Mapoly0105s0040.1) or in UNIPROT (www.uniprot.org): *Hs*TPX2 (Q9ULW0); *Gg*TPX2 (F1NW64); *Xl*TPX2 (Q6NUF4).

## Acknowledgments

We thank Momoko Nishina and Yuki Nakaoka for assistance with this project, Sebastian Hoernstein for help and advice on flow cytometry and RNA extraction, Lennard Bohlender for help with qRT-PCR, Raymundo Alfaro-Aco for comments on the TPX2 functional motifs, Peishan Yi and Mariana Costa for helpful comments on the manuscript, and the Life Imaging Center at the University of Freiburg for microscopic use. This work was funded by JSPS KAKENHI (17H06471), JSPS and DFG under the Joint Research Projects-LEAD with UKRI (to G.G.), and an Alexander von Humboldt Foundation Postdoctoral Fellowship (to E.K.). Additional support came from the German Research Foundation (DFG) under Germany’s Excellence Strategy (CIBSS – EXC-2189 – Project ID 390939984 to R.R.). The authors declare no competing interests.

## Author contributions

E. K. and G.G. designed the research project, G.G. and R.R supervised the project, E.K. and M.W.Y. performed experiments, E.K. analyzed the data, E.K., G.G. and R.R. wrote and edited the manuscript.

---

**Supplemental figure 1.**
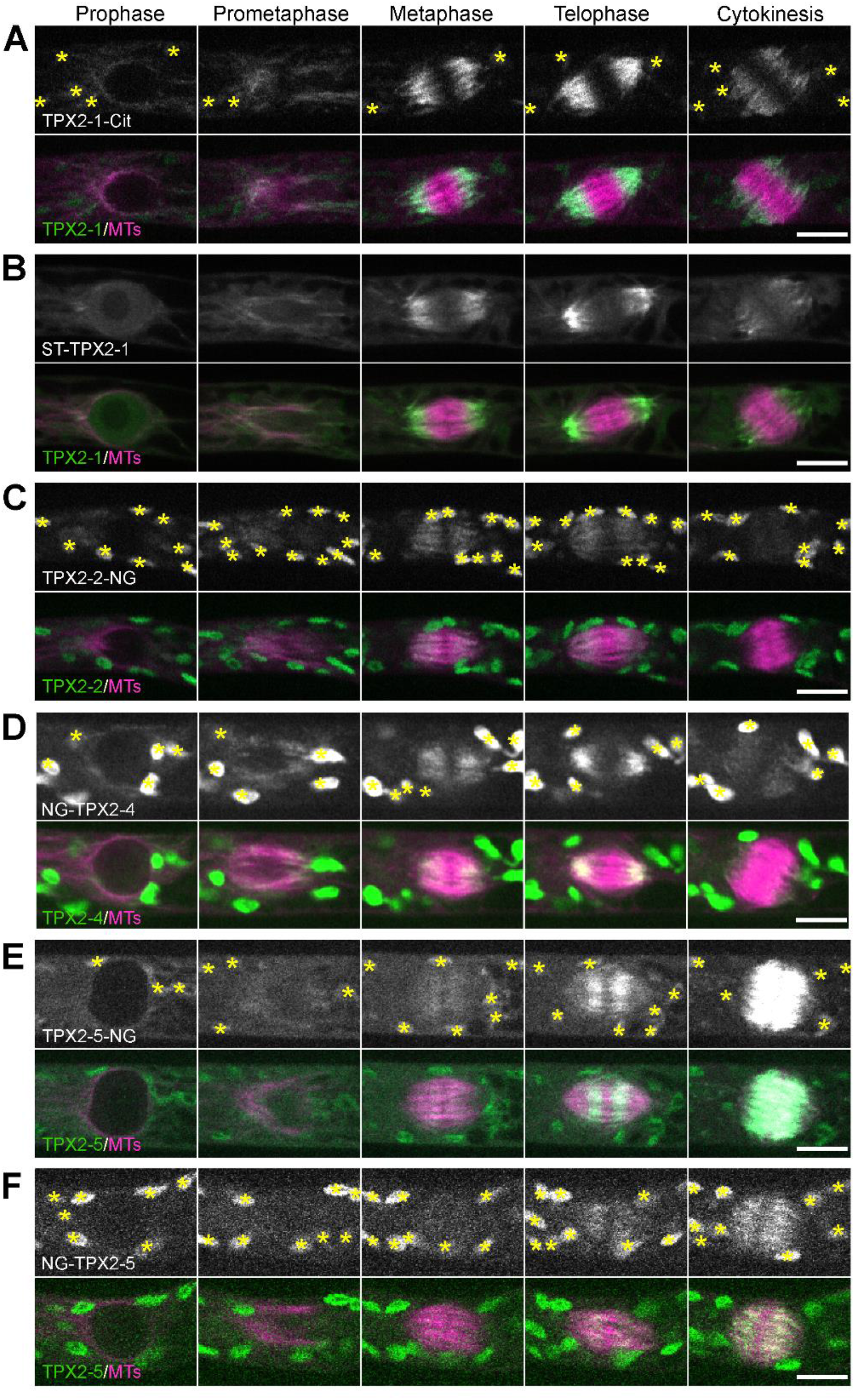
Localization of TPX2 proteins during mitosis. Live-cell imaging was performed in caulonemal apical cells of *P. patens*, expressing mCherry-tubulin and one of the following TPX2 proteins endogenously tagged with a fluorophore: (A) TPX2-1-Citrine; (B) SunTag-TPX2-1; (C) TPX2-2-mNeonGreen; (D) mNeonGreen-TPX2-4; (E) TPX2-5-mNeonGreen; (F) mNeonGreen-TPX2-5. The SunTag-TPX2-1 line also expressed scFv-GCN-sfGFP under a β-estradiol-inducible promoter. Asterisks indicate the autofluorescent chloroplasts. Bars, 10 μm.

**Supplemental figure 2.**
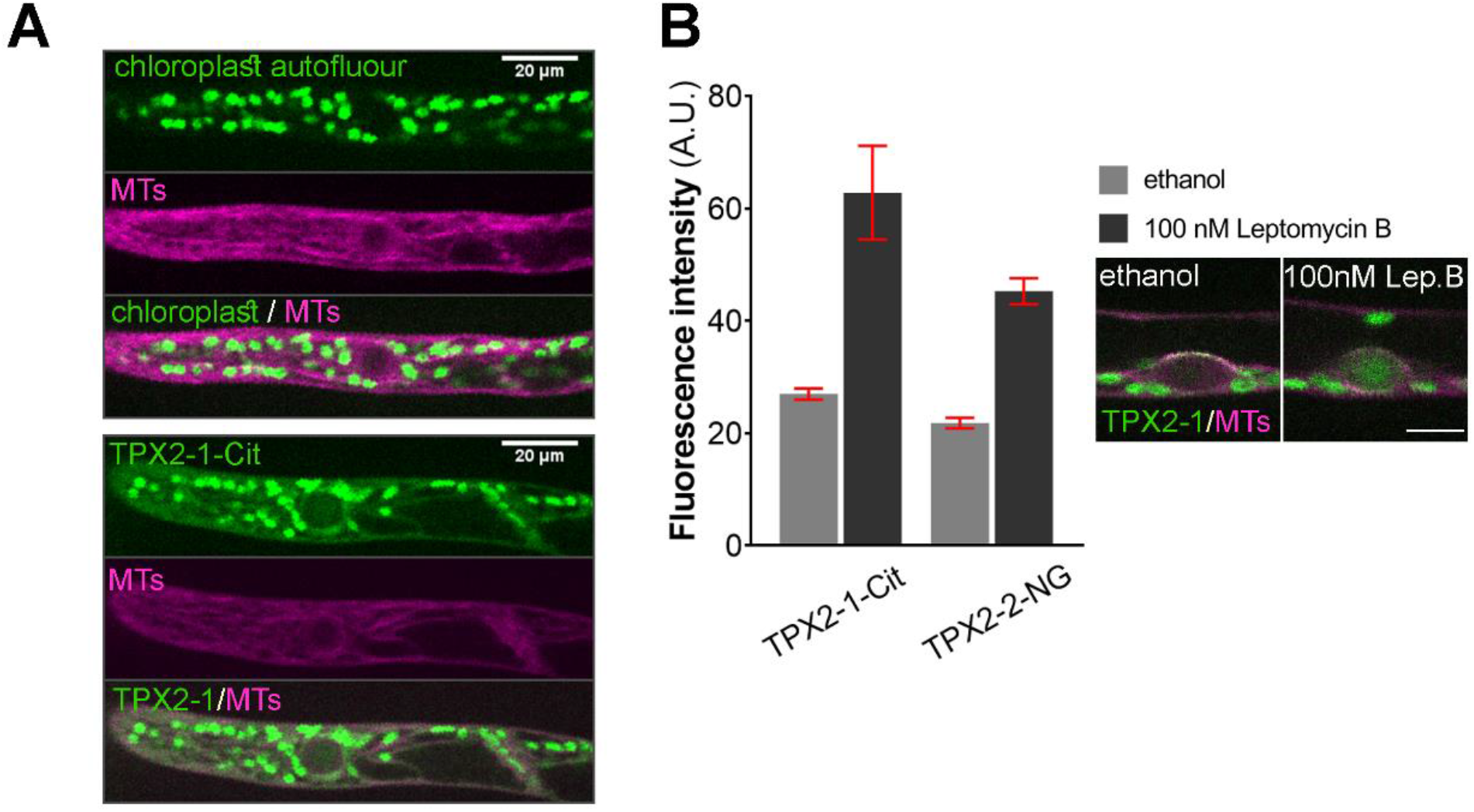
Interphase localization of TPX2. **(A)** Live-cell imaging was performed in caulonemal apical cells of *P. patens*, expressing only mCherry-tubulin (upper panels) or mCherry-tubulin and TPX2-1-Citrine (bottom panels). Cytoplasmic signals increased in the TPX2-1-Citrine line. Bars, 20 μm. **(B)** Fluorescence intensity in the nucleus before and after inhibiting nuclear export with 100 nM Leptomycin B (mean ± SEM) after subtracting the cytoplasmic background. Increase in nuclear signals after nuclear export inhibition suggests that TPX2-1-Cit and TPX2-2-NG are actively shuttled between the nucleus and cytoplasm. Bar, 10 μm.

**Supplemental figure 3.**
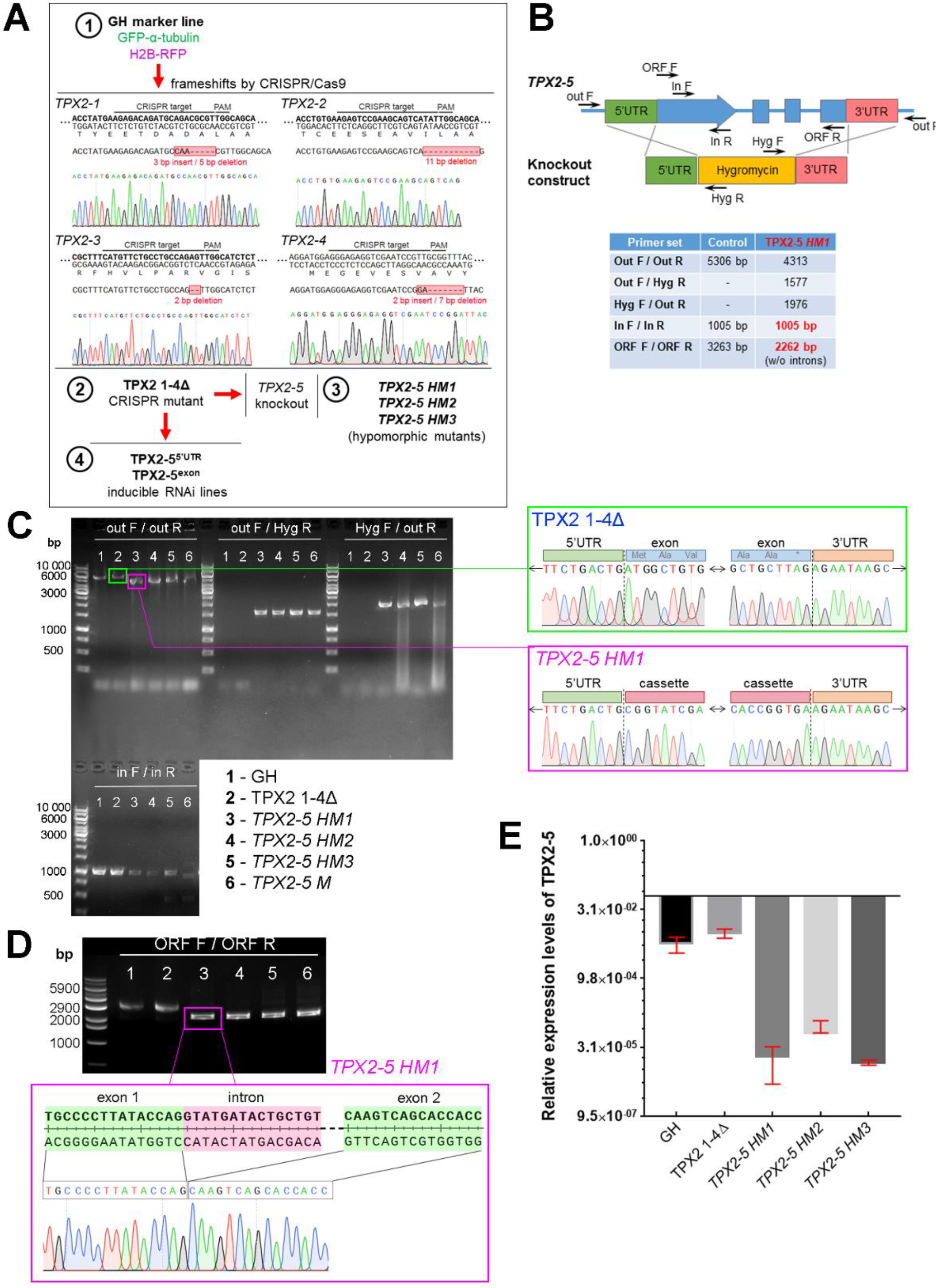
Isolation of hypomorphic *TPX2-5* mutants. **(A)** Schematic explanation of *P. patens* lines created and used in this study and representative sequencing data of frameshift mutations in the *TPX2 1-4Δ* line. **(B)** Schematic explanation of *TPX2-5* hypomorphic line selection and genotyping PCR. **(C)** Results of genotyping PCR and sequencing of the *TPX2-5* locus. **(D)** Results of amplifying full coding sequence from start to stop codons and sequencing results of the *TPX2-5* gene. Note the band shift in the *TPX2-5 HM1* line due to loss of introns. **(E)** qRT-PCR analysis of *TPX2-5* expression normalized against the expression of the internal housekeeping genes *EF1α* and *L21*. Bars represent mean values ± SD from n = 3.

**Supplemental figure 4.**
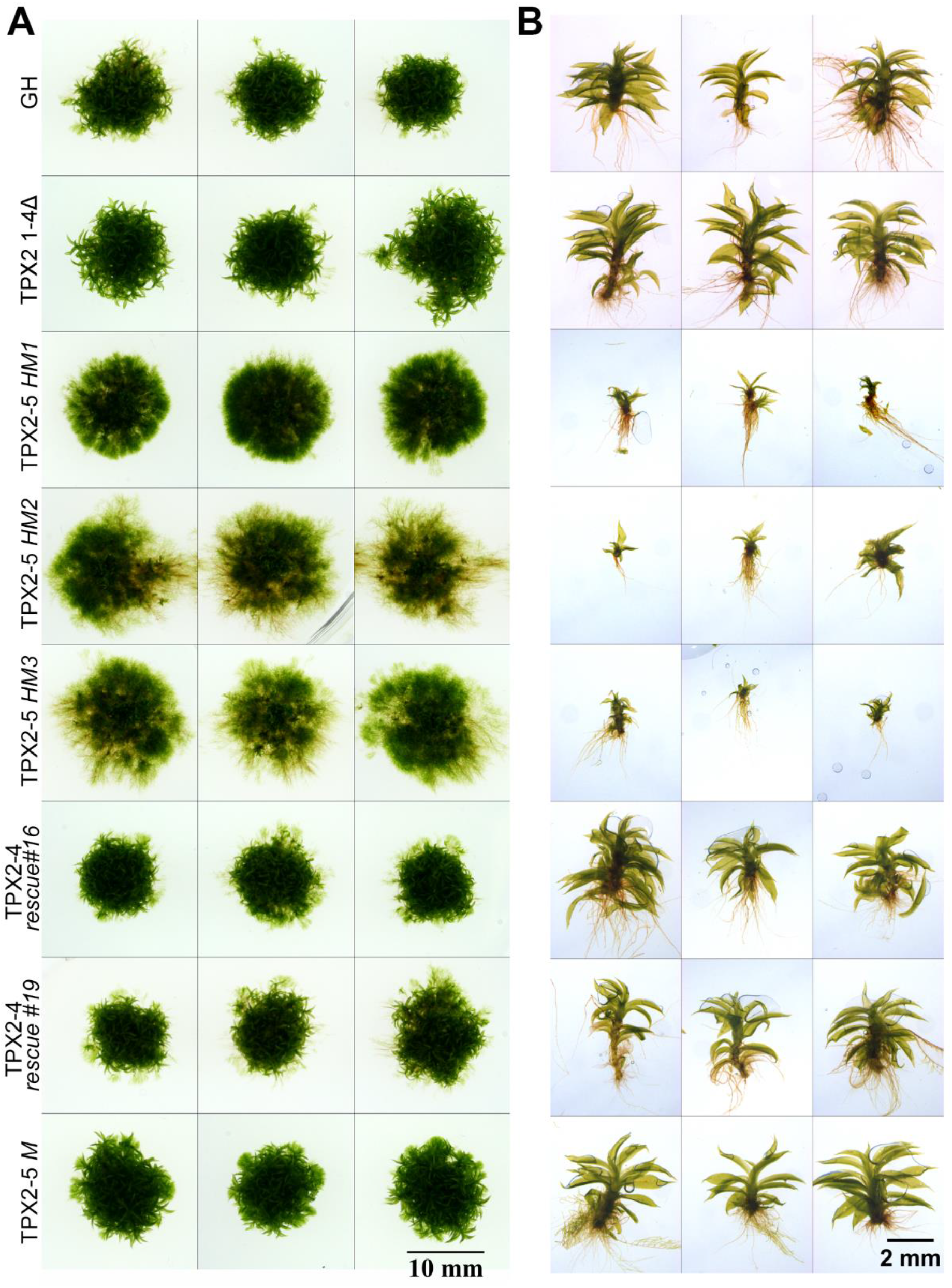
Development of *P. patens* colonies and gametophores after 4 weeks of culture. Representative images of moss colonies **(A)** or gametophores **(B)** for all lines (except RNAi) used in this study after 4 weeks of culture on BCDAT agar plates. The brightness and contrast were linearly adjusted. Some photos are the duplicates of those in Figure 2A. Bars, 10 mm and 2 mm, respectively.

**Supplemental figure 5.**
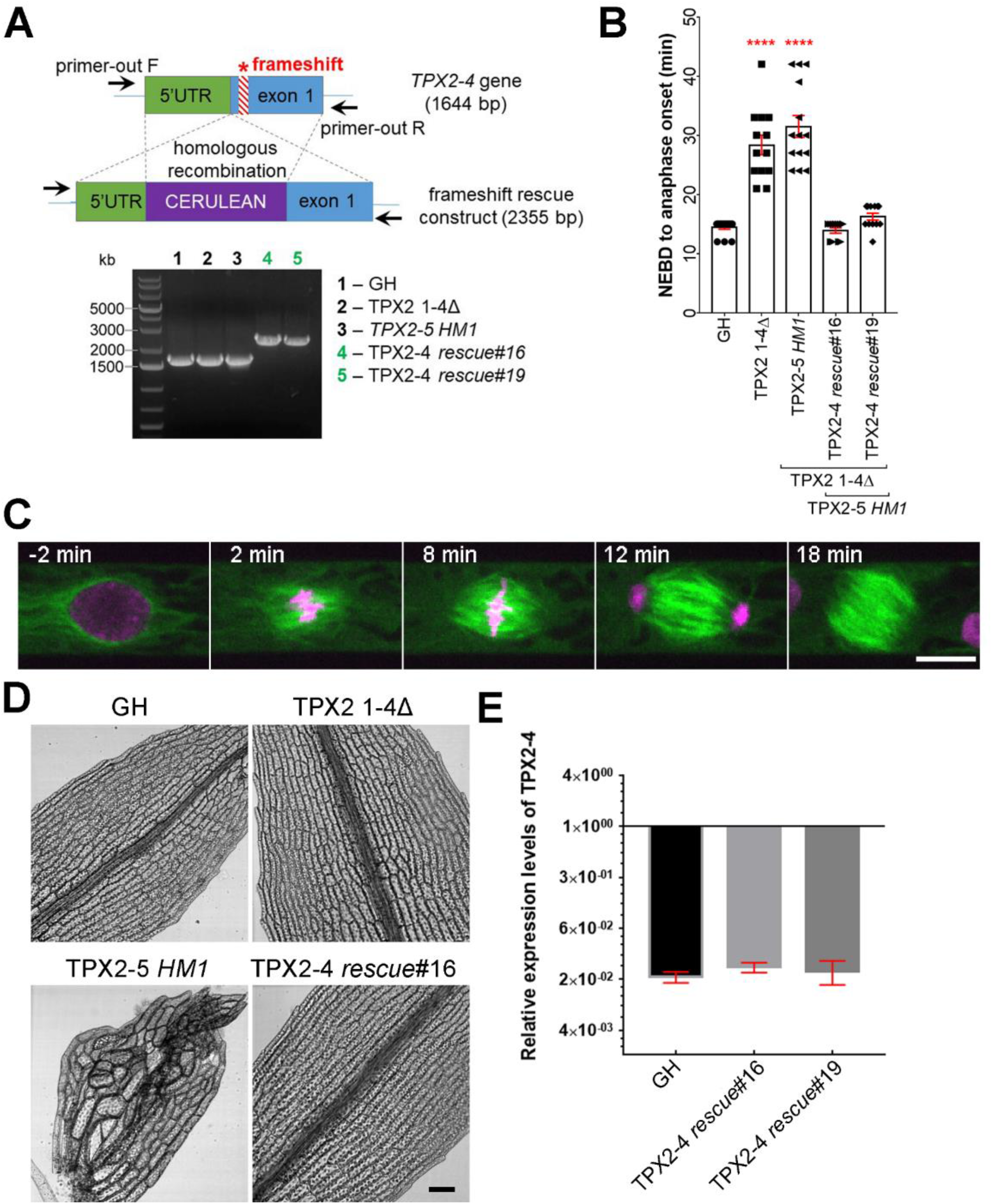
Rescue of the *TPX2-5 HM1* phenotypes by frameshift rescue of the *TPX2-4* gene. **(A)** Schematic illustration of the frameshift rescue experiment. In brief, the N-terminus-coding region of the *TPX2-4* gene in the *TPX2-5 HM1* background was tagged with Cerulean flanked with ~500 bp of the 5’-UTR and exon region (without the frameshift mutation) by homologous recombination. Construct integration was verified by PCR. **(B)** Mitotic duration of protonemal cells calculated from NEBD to anaphase onset in GH (control), *TPX2 1-4Δ, TPX2-5 HM*, and two independent *TPX2-4* rescue lines (mean ± SEM, ****p=0.0001 by one way Anova with Dunnett’s multiple comparison test against GH). **(C)** Mitotic progression in the *TPX2-4* rescue #16 line. Note that perinuclear MTs and prometaphase spindle formation were restored. Bar, 10 μm. **(D)** Representative images of gametophore leaf cells in GH, *TPX2 1-4Δ, TPX2-5 HM*, and *TPX2-4 rescue*#16 lines. Bar, 100 μm. **(E)** qRT-PCR analysis of *TPX2-4* expression normalized against the expression of the internal housekeeping genes *EF1α* and *L21*. No significant difference between GH and *TPX2-4 rescue*#16 and #19 by one-way Anova test and t-test.

**Supplemental figure 6.**
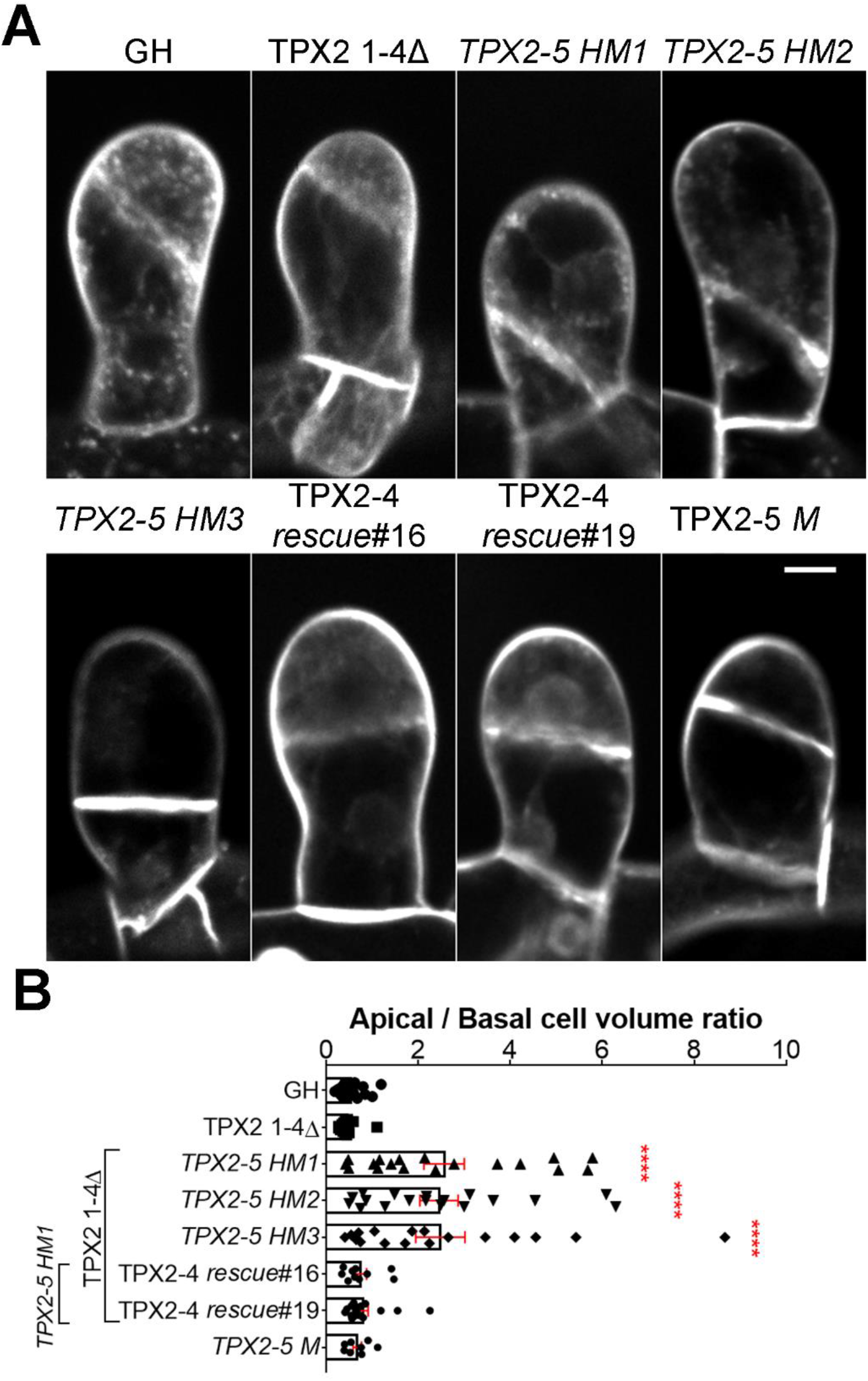
Cell division site and apical/basal cell ratio at the 2-cell stage of gametophore development in different TPX2 backgrounds. **(A)** Gametophore initial at the 2-cell stage stained with FM4-64 dye. Bar, 10 μm **(B)** The apical/basal cell volume ratio was estimated as the apical cell volume divided by the basal cell volume, measured during the 2-cell stage (mean ± SEM, ****p=0.0001 by one way ANOVA with Dunnett’s multiple comparison test against GH). n = 22, 15, 18, 18, 17, 10, 18 and 8 for GH, TPX2 1-4Δ, *TPX2-5 HM1, TPX2-5 HM2, TPX2-5 HM3*, TPX2-4 *rescue*#16, TPX2-4 *rescue*#19 and *TPX2-5 M*.

**Supplemental figure 7.**
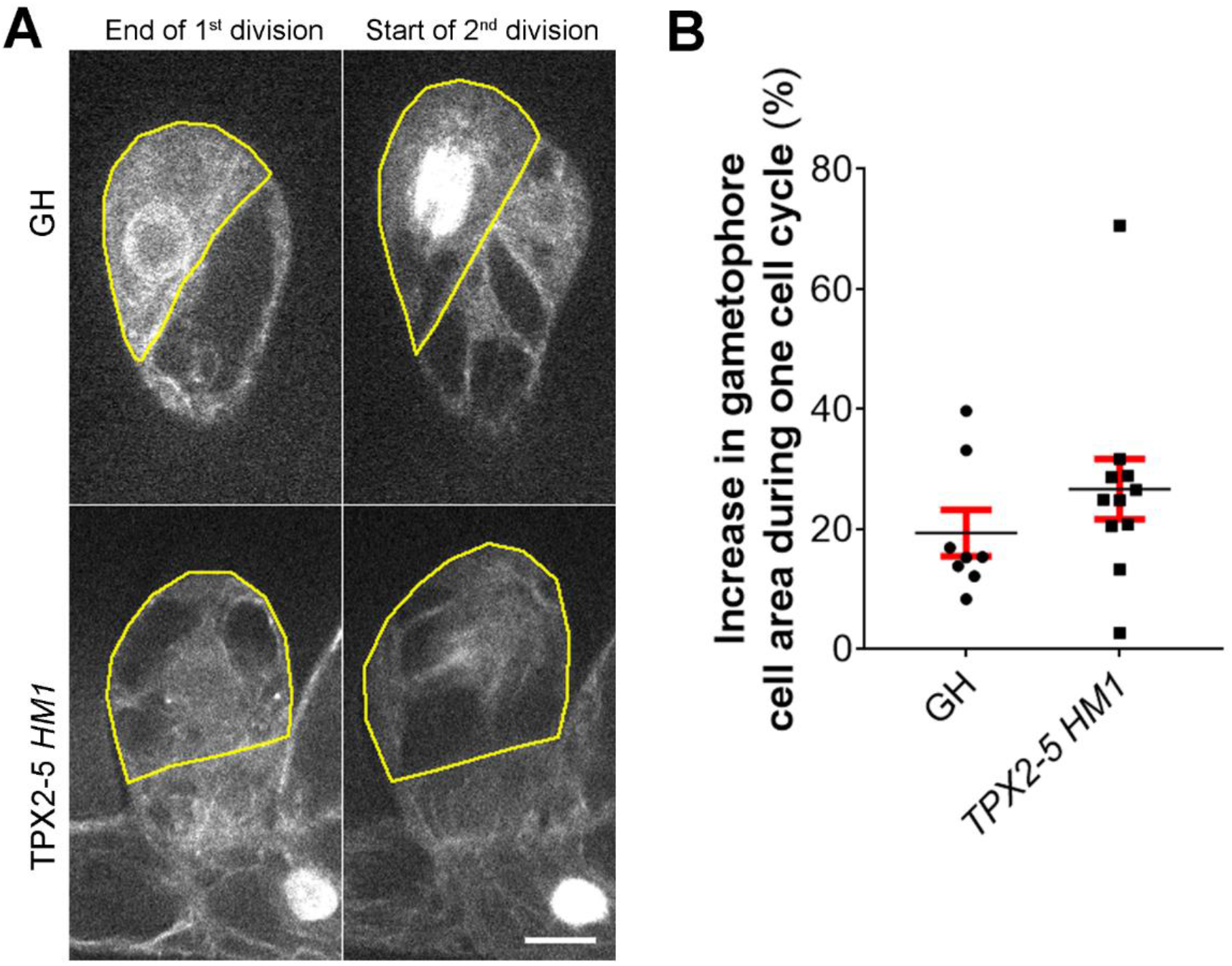
Analysis of cell growth in early gametophore development. **(A)** Representative images of cell expansion in gametophore initials during one cell cycle (from disappearance of phragmoplast to nuclear envelope breakdown) in GH and TPX2-5 HM1 lines. Cell borders are indicated with yellow lines. Images were acquired as Z-stack and the best focal plate was selected for analysis. Bar, 10 μm **(B)** Quantification of the increase in the cell area during one cell cycle. Cell area was first measured at the end of the cell division. The value was taken as 100%. The next measurement was taken at the onset of the subsequent cell division and values were plotted as increase percentage. Only gametophores with basal spindle motility were analyzed for *TPX2-5 HM1* line. Bars represent mean ± SEM.

**Supplemental figure 8.**
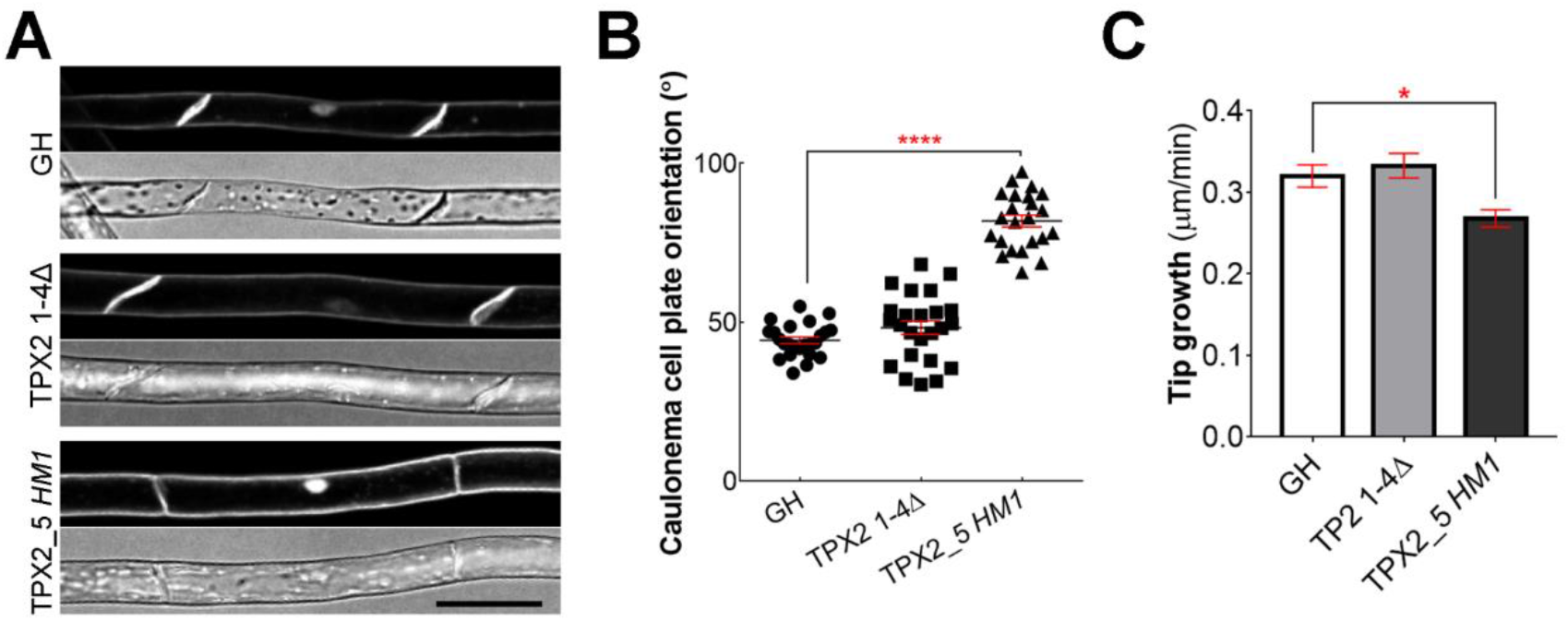
Phragmoplast orientation and tip growth defects in the caulonemal tip cells in the *TPX2-5 HM1* line. **(A)** Representative images of cell plates in caulonemal cells stained with 10 μM FM4-64 dye. Bar, 50 μm **(B)** Acute angle of caulonemal cells in GH (control), *TPX2 1-4Δ* and *TPX2_5 HM1* lines, *n* = 24, 24, and 23, respectively (mean±SEM, **** p = 0.0001 one way ANOVA with Dunnett’s multiple comparisons test) **(C)** Tip growth rate of caulonemal cells in GH, TPX2 1-4Δ and TPX2_5 *HM1* lines, *n* = 46, 50 and 48, respectively (mean±SEM, * p = 0.0144 one way ANOVA with Dunnett’s multiple comparisons test).

**Supplemental figure 9.**
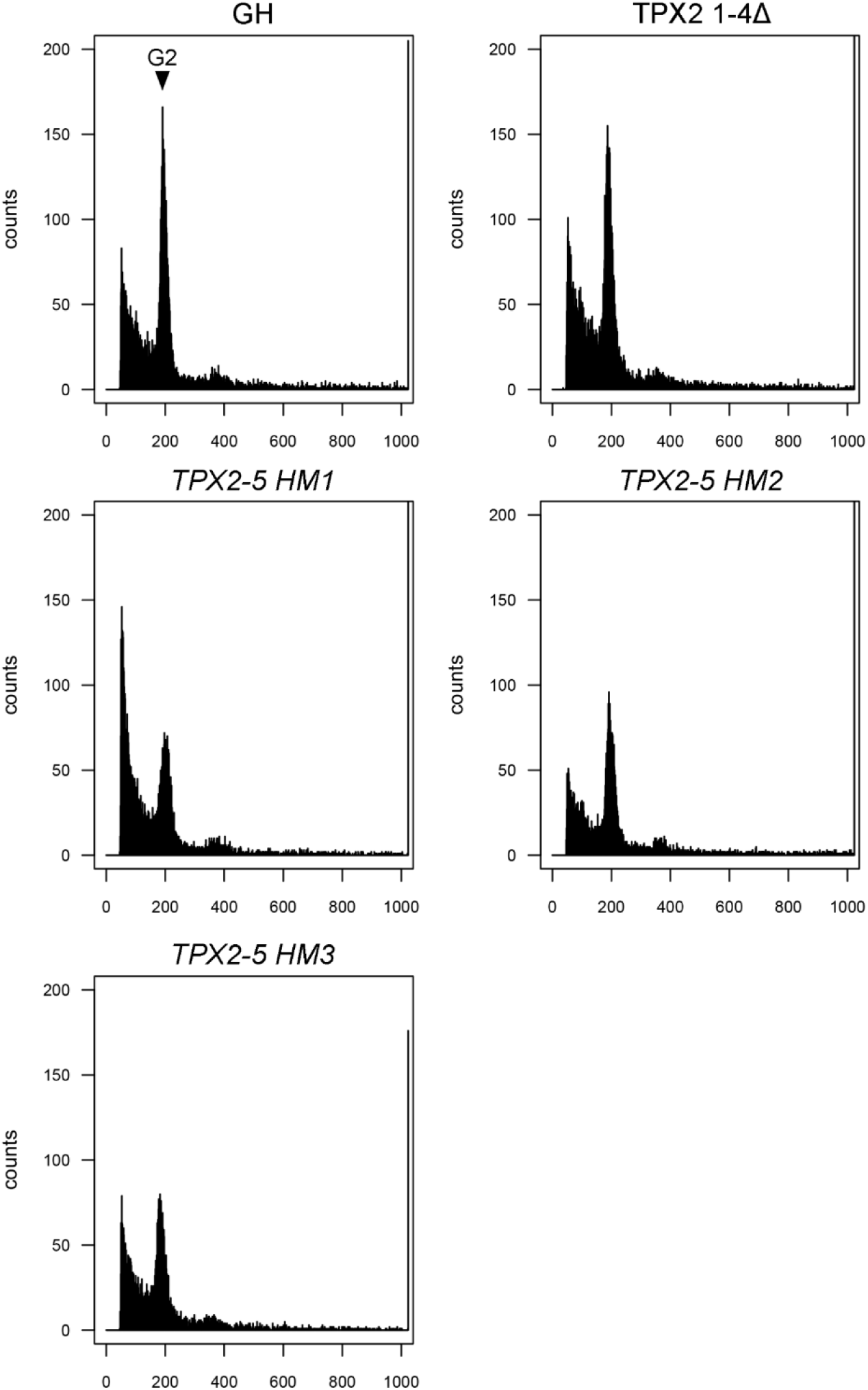
Ploidy analysis by flow cytometry. Histograms of nuclei isolated from 6-day-old protonema tissues stained with DAPI. Peak corresponding to G2 (2N) in the control line (GH) is indicated by arrow.

**Video 1. Localization of TPX2 proteins during mitosis**

Live-cell imaging was performed in *P. patens* apical caulonemal cells expressing mCherry-tubulin (magenta) and one of the following tagged proteins (green): TPX2-1-Citrine, TPX2-2-mNeonGreen, mNeonGreen-TPX2-4, or TPX2-5-mNeonGreen. Images were acquired every 30 s in a single focal plane. Bar, 10 μm.

**Video 2. 3D projection of gametophore initial cells after the first division.** Gametophore initials were stained with 10 μM FM4-64. Cell segmentation was done with PlantSeg. The apical cell of the gametophore initial is shown in pink, while the basal cell in green. Bar, 20 μm.

**Video 3. Spindle motility underlies the erroneous phragmoplast positioning in *TPX2-5 HM1* gametophore initial cells**

Live-cell imaging was performed in *P. patens* gametophore initial cells expressing mCherry-tubulin and histone H2B-mCherry (chromosomes and MTs are labeled with the same color). Images were acquired as a z-stack (20 μm in 2.5 μm steps) every 5 min, and the best focal plane was presented. Bar, 10 μm.

**Video 4. Spindle positioning defects in later gametophore development in the *TPX2-5 HM1* mutant.**

Representative videos of spindle misorientation and/or motility in *TPX2-5 HM1* gametophore initial cells during second (left cell) or later (right cell) cell divisions. The spindle axis is indicated by a cyan line. Images were acquired as a z-stack (20 μm, 2.5 μm step) every 5 min, and the best focal plane was presented. Bar, 10 μm.

**Video 5. Spindle-collapse phenotype in the gametophore initial of *TPX2-5 HM1* mutants**

Representative video of spindle collapse followed by chromosome missegregation, observed in approximately 10% of *TPX2-5 HM1* gametophore initial cells. Images were acquired as a z-stack (20 μm, 2.5 μm step) every 5 min, and the best focal plane was presented. Bar, 10 μm.

**Video 6. Mitotic defects in the *TPX2-5* RNAi lines**

Representative images of mitotic defects in the *TPX2-5* RNAi lines. Live-cell imaging was performed in *P. patens* protonemal apical cells expressing GFP-tubulin (green) and histone H2B-mCherry (magenta). Images were acquired at a single focal plane every 2 min. Bar, 10 μm.

**Supplementary table 1. Transgenic *Physcomitrium patens* lines used in this study**

**Supplementary table 2. Plasmids and primers used in this study**

## References

Alfaro-Aco R, Thawani A, Petry S. 2017. Structural analysis of the role of TPX2 in branching microtubule nucleation. J Cell Biol 177:7–11. doi:10.1083/jcb.200611141

Almonacid M, Terret MÉ, Verlhac MH. 2014. Actin-based spindle positioning: New insights from female gametes. J Cell Sci 127:477–483. doi:10.1242/jcs.142711

Ambrose JC, Cyr R. 2008. Mitotic spindle organization by the preprophase band. Mol Plant 1:950–960. doi:10.1093/mp/ssn054

Arima K, Tamaoki D, Mineyuki Y, Yasuhara H, Nakai T, Shimmen T, Yoshihisa T, Sonobe S. 2018. Displacement of the mitotic apparatuses by centrifugation reveals cortical actin organization during cytokinesis in cultured tobacco BY-2 cells. J Plant Res 131:803–815. doi:10.1007/s10265-018-1047-4

Bergstralh DT, Dawney NS, Johnston DS. 2017. Spindle orientation : a question of complex positioning. Development 5:1137–1145. doi:10.1242/dev.140764

Boruc J, Deng X, Mylle E, Besbrugge N, Durme M Van, Demidov D, Tomaštíková ED, Tan T-RC, Vandorpe M, Eeckhout D, Beeckman T, Nowack M, Jaeger G De, Lin H, Liu B, Damme D Van. 2019. The TPX2-LIKE PROTEIN 3 is the primary activator of α Aurora kinases and is essential for embryogenesis. Plant Physiol. doi:10.1101/466276

Brejšková L, Hála M, Rawat A, Soukupová H, Cvrčková F, Charlot F, Nogué F, Haluška S, Žárský V. 2021. SEC6 exocyst subunit contributes to multiple steps of growth and development of Physcomitrella (Physcomitrium patens). Plant J. doi:10.1111/tpj.15205

Brownfield L, Yi J, Jiang H, Minina EA, Twell D, Köhler C. 2015. Organelles maintain spindle position in plant meiosis. Nat Commun 6. doi:10.1038/ncomms7492

Buschmann H, Müller S. 2019. Update on plant cytokinesis: rule and divide. Curr Opin Plant Biol 52:97–105. doi:10.1016/j.pbi.2019.07.003

Doonan JH, Cove DJ, Corke FMK, Lloyd CW. 1987. Pre-prophase band of microtubules, absent from tip-growing moss filaments, arises in leafy shoots during transition to intercalary growth. Cell Motil Cytoskeleton 7:138–153. doi:10.1002/cm.970070206

Doonan JH, Cove DJ, Lloyd CW. 1988. Microtubules and microfilaments in tip growth : evidence that microtubules impose polarity on protonemal growth in Physcomitrella patens. J Cell Sci 89:533–540.

Duan X, Li Y, Yi K, Guo F, Wang HY, Wu PH, Yang J, Mair DB, Morales EA, Kalab P, Wirtz D, Sun SX, Li R. 2020. Dynamic organelle distribution initiates actin-based spindle migration in mouse oocytes. Nat Commun 11:277. doi:10.1038/s41467-019-14068-3

Jordan MANN, Toso RJ, Thrower D, Wilson L. 1993. Mechanism of Mitotic Block and Inhibition of Cell Proliferation by Taxol at Low Concentrations. Proc Natl Acad Sci 90:9552–9556.

Kimata Y, Kato T, Higaki T, Kurihara D, Yamada T, Segami S, Morita MT, Maeshima M, Hasezawa S, Higashiyama T, Tasaka M, Ueda M. 2019. Polar vacuolar distribution is essential for accurate asymmetric division of Arabidopsis zygotes. Proc Natl Acad Sci U S A 116:2338–2343. doi:10.1073/pnas.1814160116

Kiyomitsu T. 2019. The cortical force-generating machinery: how cortical spindle-pulling forces are generated. Curr Opin Cell Biol 60:1–8. doi:10.1016/j.ceb.2019.03.001

Kosetsu K, Murata T, Yamada M, Nishina M, Boruc J, Hasebe M, Van Damme D, Goshima G. 2017. Cytoplasmic MTOCs control spindle orientation for asymmetric cell division in plants. Proc Natl Acad Sci 114:E8847–E8854. doi: 10.1073/pnas.1713925114

Kozgunova E, Nishina M, Goshima G. 2019. Kinetochore protein depletion underlies cytokinesis failure and somatic polyploidization in the moss Physcomitrella patens. Elife 8:1–16. doi:10.7554/eLife.43652

Leong SY, Yamada M, Yanagisawa N, Goshima G. 2018. SPIRAL2 Stabilises Endoplasmic Microtubule Minus Ends in the Moss *Physcomitrella patens*. Cell Struct Funct 43:53–60. doi:10.1247/csf.18001

Lipka E, Gadeyne A, Stöckle D, Zimmermann S, De Jaeger G, Ehrhardt DW, Kirik V, Van Damme D, Müller S. 2014. The phragmoplast-orienting kinesin-12 class proteins translate the positional information of the preprophase band to establish the cortical division zone in Arabidopsis thaliana. Plant Cell 26:2617–2632. doi:10.1105/tpc.114.124933

Livak KJ, Schmittgen TD. 2001. Analysis of relative gene expression data using real-time quantitative PCR and the 2-ΔΔCT method. Methods 25:402–408. doi:10.1006/meth.2001.1262

Livanos P, Müller S. 2019. Division Plane Establishment and Cytokinesis. Annu Rev Plant Biol 70:239–267. doi:10.1146/annurev-arplant-050718-100444

Lopez-Obando M, Hoffmann B, Gery C, Guyon-Debast A, Teoule E, Rameau C, Bonhomme S, Nogue F. 2016. Simple and Efficient Targeting of Multiple Genes Through CRISPR-Cas9 in Physcomitrella patens. G3 Genes|Genomes|Genetics 6:3647–3653. doi:10.1534/g3.116.033266

Martinez P, Luo A, Sylvester A, Rasmussen CG. 2017. Proper division plane orientation and mitotic progression together allow normal growth of maize. Proc Natl Acad Sci U S A 114:2759–2764. doi:10.1073/pnas.1619252114

Miki T, Nakaoka Y, Goshima G. 2016. Live Cell Microscopy-Based RNAi Screening in the Moss Physcomitrella patens. Methods Mol Biol 1470:225–246. doi:10.1007/978-1-4939-6337-9_18

Moody LA, Kelly S, Rabbinowitsch E, Langdale JA. 2018. Genetic Regulation of the 2D to 3D Growth Transition in the Moss Physcomitrella patens. Curr Biol 28:473–478.e5. doi:10.1016/j.cub.2017.12.052

Müller S. 2019. Plant cell division — defining and finding the sweet spot for cell plate insertion. Curr Opin Cell Biol 60:9–18. doi:10.1016/j.ceb.2019.03.006

Nakaoka Y, Miki T, Fujioka R, Uehara R, Tomioka A, Obuse C, Kubo M, Hiwatashi Y, Goshima G. 2012. An Inducible RNA Interference System in Physcomitrella patens Reveals a Dominant Role of Augmin in Phragmoplast Microtubule Generation. Plant Cell 24:1478–1493. doi: 10.1105/tpc.112.098509

Ôta T. 1961. The Role of Cytoplasm in Cytokinesis of Plant Cells. Cytologia (Tokyo) 26:428–447. doi:10.1508/cytologia.26.428

Petry S. 2016. Mechanisms of Mitotic Spindle Assembly. Annu Rev Biochem 85:659–683. doi:10.1146/annurev-biochem-060815-014528

Petry S, Groen AC, Ishihara K, Mitchison TJ, Vale RD. 2013. Branching microtubule nucleation in xenopus egg extracts mediated by augmin and TPX2. Cell 152:768–777. doi:10.1016/j.cell.2012.12.044

Rasmussen CG, Bellinger M. 2018. An overview of plant division-plane orientation. New Phytol 219:505–512. doi:10.1111/nph.15183

Schaefer E, Belcram K, Uyttewaal M, Duroc Y, Goussot M, Legland D, Laruelle E, De Tauzia-Moreau ML, Pastuglia M, Bouchez D. 2017. The preprophase band of microtubules controls the robustness of division orientation in plants. Science (80-) 356:186–189. doi:10.1126/science.aal3016

Schween G, Gorr G, Hohe A, Reski R. 2003. Unique tissue-specific cell cycle in Physcomitrella. Plant Biol 5:50–58. doi:10.1055/s-2003-37984

Schween G, Schulte J, Reski R. 2005. Effect of Ploidy Level on Growth, Differentiation, and Morphology in Physcomitrella patens. Bryologist 108:27–35.

Smertenko A, Assaad F, Baluška F, Bezanilla M, Buschmann H, Drakakaki G, Hauser MT, Janson M, Mineyuki Y, Moore I, Müller S, Murata T, Otegui MS, Panteris E, Rasmussen C, Schmit AC, Šamaj J, Samuels L, Staehelin LA, Van Damme D, Wasteneys G, Žárský V. 2017. Plant Cytokinesis: Terminology for Structures and Processes. Trends Cell Biol 27:885–894. doi:10.1016/j.tcb.2017.08.008

Tomaštíková E, Demidov D, Jeřábková H, Binarová P, Houben A, Doležel J, Petrovská B. 2015. TPX2 Protein of Arabidopsis Activates Aurora Kinase 1, But Not Aurora Kinase 3 In Vitro. Plant Mol Biol Report 33:1988–1995. doi:10.1007/s11105-015-0890-x

Vargas-Hurtado D, Brault JB, Piolot T, Leconte L, Da Silva N, Pennetier C, Baffet A, Marthiens V, Basto R. 2019. Differences in Mitotic Spindle Architecture in Mammalian Neural Stem Cells Influence Mitotic Accuracy during Brain Development. Curr Biol 29:2993–3005.e9. doi:10.1016/j.cub.2019.07.061

Verma DPS. 2001. Cytokinesis and building the cell wall in plants. Annu rev plant Physiol 52:751–784.

Vos JW, Pieuchot L, Evrard J-L, Janski N, Bergdoll M, De Ronde D, Pérez LH, Sardon T, Vernos I, Schmit A-C. 2008. The Plant TPX2 Protein Regulates Prospindle Assembly before Nuclear Envelope Breakdown. Plant Cell 20:2783–2797. doi:10.1105/tpc.107.056796

Wolny A, Cerrone L, Vijayan A, Tofanelli R, Barro AV, Louveaux M, Wenzl C, Strauss S, Wilson-Sánchez D, Lymbouridou R, Steigleder SS, Pape C, Bailoni A, Duran-Nebreda S, Bassel G, Lohmann JU, Tsiantis M, Hamprecht FA, Schneitz K, Maizel A, Kreshuk A. 2020. Accurate and versatile 3D segmentation of plant tissues at cellular resolution. Elife 9:1–34. doi:10.7554/eLife.57613

Yamada M, Goshima G. 2018. The KCH kinesin drives nuclear transport and cytoskeletal coalescence to promote tip cell growth in Physcomitrella patens. Plant Cell 30:1496–1510. doi:10.1105/tpc.18.00038

Yamada M, Miki T, Goshima G. 2016. Imaging Mitosis in the Moss Physcomitrella patens. Methods Mol Biol 1413:293–326. doi:10.1007/978-3-319-02904-7

Yoneda A, Akatsuka M, Hoshino H, Kumagai F, Hasezawa S. 2005. Decision of spindle poles and division plane by double preprophase bands in a BY-2 cell line expressing GFP-tubulin. Plant Cell Physiol 46:531–538. doi:10.1093/pcp/pci055

